# Secretion of redox-active metabolites as a general strategy for iron acquisition in plants

**DOI:** 10.1101/168104

**Authors:** Jakub Rajniak, Ricardo F. Giehl, Evelyn Chang, Irene Murgia, Nicolaus von Wirén, Elizabeth S. Sattely

## Abstract

Iron is an essential but often poorly bioavailable nutrient due to its low solubility, especially in alkaline soils. Plants have evolved at least two distinct strategies to extract iron from soil: solubilization of ferric iron by phytosiderophores, and reduction to the ferrous form at the root surface followed by direct import. Here, we describe the discovery of a novel redox-active catecholic metabolite, termed sideretin, which derives from the coumarin fraxetin, and is the primary molecule exuded by *Arabidopsis thaliana* roots in response to iron deficiency. Using a combination of metabolomics, heterologous expression, and coexpression analysis we have identified two enzymes that complete the biosynthetic pathway of sideretin. Chemical characterization of synthetic sideretin and biological assays with pathway mutants suggest that sideretin is critical for iron nutrition and support a role for small molecule-mediated iron reduction in *A. thaliana* under iron-limiting growth conditions. Further, we show that sideretin production is evolutionarily ancient and occurs in eudicot species only distantly related to *A. thaliana*. In addition to sideretin, untargeted metabolomics of the root exudates of various eudicots revealed the production of structurally diverse redox-active molecules in response to iron deficiency. Our results indicate that secretion of small molecule reductants by roots may be a widespread and previously underappreciated component of the reduction-based iron uptake strategy.

## Introduction

Iron (Fe) is a key micronutrient required for the growth of all organisms, fulfilling numerous, diverse biochemical roles. In plants, iron is essential for various cellular processes, such as respiration and chlorophyll biosynthesis, and serves as a cofactor for many enzymes involved in electron or oxygen transfer processes, such as cytochrome P450s (CYPs) and 2-oxoglutarate-dependent dioxygenases (2-ODDs)^1,2^.

While nominally abundant in the Earth’s crust, iron is only poorly bioavailable due to the very low solubility product of the ferric form Fe(III) with several common soil anions, such as hydroxide and phosphate. Even though the ferrous form Fe(II) is far more soluble, it is readily oxidized to Fe(III) in well-aerated soils. To cope with poor iron availability, land plants have evolved at least two distinct strategies for iron mobilization and uptake^3,4^. The chelation-based mechanism, also known as Strategy II, is confined to true grasses (family Poaceae) and relies on the exudation^5^ and re-uptake of mugineic acid-type phytosiderophores^6,7^, which solubilize and bind Fe(III)^8^. All other plant families are thought to employ a reduction-based mechanism (Strategy I), which involves acidification of the rhizosphere via proton secretion^9^, enzymatic reduction of iron chelates at the root surface^10^, and secondary active transport of Fe(II) across the plasma membrane^11,12^. Notably, efficient root-mediated Fe(III) reduction in Strategy I plants depends on the presence of soluble Fe(III)-chelates derived, e.g., from the soil^13^. Several enzymes of the phenylpropanoid pathway are upregulated under iron deficiency in *Arabidopsis*^14,15^, suggesting that low molecular-weight compounds released from Strategy I plants are also involved in mobilization of iron from insoluble pools in the soil or root apoplast^16,17^; however, this observation has not been explained mechanistically.

Recently, several groups have independently reported that root exudation of coumarins derived from the phenylpropanoid pathway are required for efficient iron uptake, especially in alkaline conditions^15,18–20^. The dedicated oxoglutarate-dependent oxidase F6’H1^15,19–21^ and the ABC-type transporter PDR9^18^ have been shown to be involved in coumarin biosynthesis and export, respectively. In line with a role for coumarins in Fe acquisition, both *f6’h1* and *pdr9* mutants show severe symptoms of iron deficiency relative to WT plants when iron was supplied in an insoluble form^15,18–20^. Coumarins bearing a catechol motif can potentially contribute to Fe acquisition by two mechanisms: (1) solubilization of ferric Fe precipitates since catechols are potent Fe(III) ligands, and (2) reduction of Fe(III) to Fe(II) for direct import^22^. However, in reports to date, the two known natural catecholic coumarins found in the exudates of iron-deficient wild-type plants, esculetin and fraxetin, were produced at insufficient levels to explain the Fe deficiency of *f6’h1* and *pdr9* plants, while scopoletin, lacking a catechol motif, is incapable of iron chelate formation or reduction^18,19^. It has been suggested that scopoletin may be further oxidized to catecholic coumarins, perhaps in the rhizosphere^18,19^.

Here, we describe the isolation, chemical synthesis, biosynthesis, and biological role of a novel catecholic coumarin which we have termed sideretin (5,7,8-trihydroxy-6-methoxycoumarin), produced by *A. thaliana*. We show that sideretin is the major coumarin exuded into the rhizosphere under iron-deficient conditions, while structurally related coumarins such as scopoletin, fraxetin and esculetin are exuded at lower levels. Using a combination of transcriptome analysis, targeted metabolomics of transfer-DNA insertion mutants, *in vitro* chemical characterization, and heterologous pathway reconstitution, we show that sideretin is biosynthesized from scopoletin via two successive hydroxylations catalyzed by a 2-ODD (S8H; At3g12900) and a CYP enzyme (CYP82C4; At4g31940). Sideretin can efficiently mobilize and reduce insoluble Fe(III) and rescue chlorotic phenotypes of various biosynthetic mutants and wild-type plants grown under conditions of low iron availability. We also show that the sideretin biosynthetic pathway arose early during the evolution of flowering plants, but appears to have been sporadically lost in various lineages. Interestingly, several Strategy I plants that do not produce sideretin exude distinct redox-active molecules under iron deficiency. Together, these data indicate that exudation of various redox-active small molecules into the rhizosphere is a key component of iron acquisition in plants.

## Results

### Sideretin biosynthetic pathway elucidation

In order to obtain a comprehensive picture of the metabolic changes that occur in *A. thaliana* plants under iron deficiency, we carried out comparative metabolomic studies of Columbia-0 (Col-0) ecotype seedlings grown hydroponically under iron-abundant and iron-depleted conditions at pH 5.7. In our setup, seeds were germinated on a buoyant PTFE mesh floating on liquid medium. This cultivation method allowed for axenic growth, easy separation of root and aerial tissue, and collection of spent medium without potentially injuring the roots, which could lead to spurious observation of naturally root-confined metabolites in the medium. Comparative analysis of spent medium methanolic extracts of 12 day-old Col-0 seedlings by HPLC-ESI-MS revealed two UV-active peaks (**1** and **2**) that strongly accumulated under iron-depleted conditions (Fig. 1a). The corresponding m/z [M+H]^+^ values of 223.0237 and 225.0394 are consistent with neutral compound molecular formulas of C_10_H_6_O_6_ (**1**) and C_10_H_8_O_6_ (**2**), respectively. Compound **2** differs from scopoletin, the primary coumarin present in *A. thaliana* roots, by two additional hydroxylations, while **1** is the dehydrogenated counterpart of **2**. We observed only trace quantities of the known coumarins scopoletin, fraxetin, and esculetin in Col-0 exudates (Supplementary Fig. 1a). We note that m/z = 225.0394 has been detected as a highly-induced signal in root exudates under iron deficiency in previous MS-based metabolomic studies^18–20^, but neither the structure nor the significance of this metabolite has been elucidated. In root tissues grown under iron-depleted conditions, we primarily observed accumulation of mass signals corresponding to the glucosides of **1** (Supplementary Figs. 1b and 2), while no significant metabolic differences were found between iron-sufficient and iron-deficient aerial tissues using our reverse-phase liquid chromatography analysis method (data not shown).

**Figure 1.**
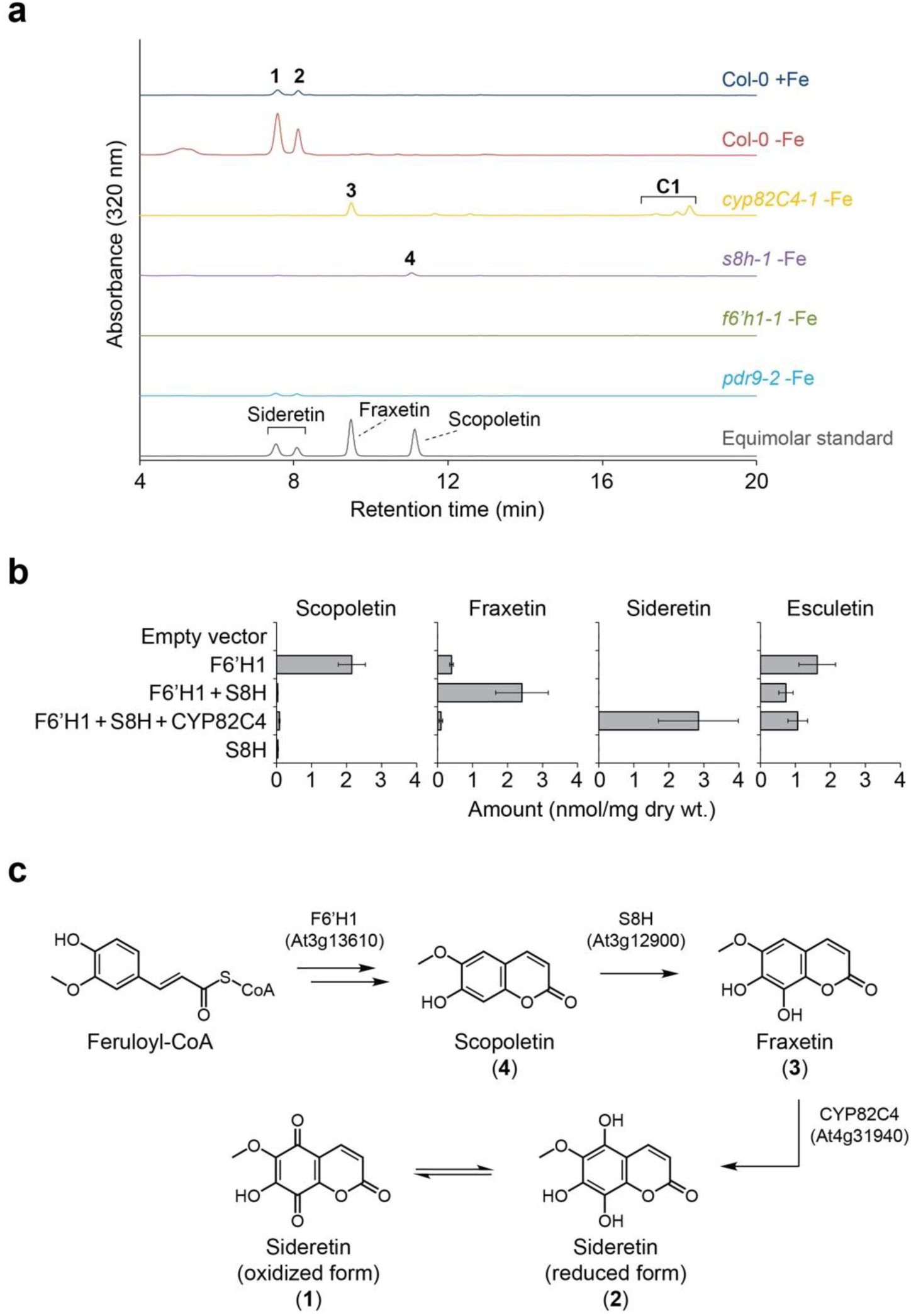
Metabolomics of T-DNA insertion lines and heterologous gene expression reveal the sideretin biosynthetic pathway in *A. thaliana*. (a) Representative UV-Vis absorbance traces for spent medium extracts of hydroponically grown *A. thaliana* lines under Fe deficiency at pH = 5.7; data are representative of three biological replicates. Peak labels correspond to compounds shown in panel c. (b) Levels of oxidized coumarins in glucosidase-treated extracts of *N. benthamiana* leaves transiently expressing subsets of sideretin pathway enzymes; data represent mean ± s.d. of four biological replicates. (c) Schematic of sideretin biosynthesis in *A. thaliana*.

We postulated that compound **2** represents a doubly hydroxylated analog of scopoletin with a catechol moiety, and **1** is the corresponding quinone. Since we could not easily obtain enough of these compounds for structural characterization directly from spent medium, we chose instead to profile various transfer-DNA (T-DNA) insertion lines potentially deficient in their biosynthesis to gain clues about structure. First, we profiled knockout lines of feruloyl-CoA 6’-hydroxylase 1 (F6’H1), the enzyme responsible for scopoletin biosynthesis in *A. thaliana*^21^. A complete absence of both compounds as well as other prominent UV-active peaks in medium of *f6’h1* lines grown under iron-depleted conditions demonstrates that they are indeed scopoletin-derived (Fig. 1a). Next, we sought to identify biosynthetic enzymes responsible for the downstream oxidation of scopoletin. One strong candidate is *CYP82C4* (*At4g31940*), which encodes a cytochrome P450 enzyme, whose expression is highly upregulated by iron deficiency and strongly correlates to the Fe(II) root importer *IRT1* under various abiotic stress conditions (Supplementary Fig. 3) ^23^. Furthermore, CYP82C4 has been shown to hydroxylate 8-methoxypsoralen, a furanocoumarin structurally similar to scopoletin^24^. We therefore obtained two independent *cyp82C4* T-DNA insertion lines for profiling. Quantitative PCR (qPCR) analysis confirmed that *CYP82C4* expression in roots is strongly induced under iron deficiency, but abolished in these T-DNA lines (Supplementary Fig. 4a). Metabolomic profiling under iron-depleted conditions further showed that both **1** and **2** are absent from root exudates of these lines. However, unlike for *f6’h1*, another prominent UV-active peak (**3**) was observed in *cyp82C4* spent medium, with m/z [M+H]^+^ = 209.0444, corresponding to C_10_H_8_O_5_. When compared with an authentic standard, this compound was identified as fraxetin (8-hydroxyscopoletin) (Fig. 1a). Therefore, we suspected that fraxetin is the direct precursor leading to **1** and **2**. To further test this hypothesis, we expressed CYP82C4 in yeast and incubated the microsomal fraction with fraxetin and NADPH. As expected, we observed *in vitro* formation of compounds identical to Col-0-derived **1** and **2** (Supplementary Fig. 5a). Taken together, these data indicate that CYP82C4 is responsible for the final step in the biosynthetic pathway leading to **1** and **2** from scopoletin. Interestingly, *cyp82C4* spent medium contains additional UV-active peaks (labeled **C1** in Fig. 1a) that correspond to non-specific cycloaddition products of fraxetin with coniferyl and sinapyl alcohols (also referred to as “cleomiscosins”^25^; Supplementary Fig. 6).

Based on the known activity of CYP82C4 as an 8-methoxypsoralen 5-hydroxylase^24^, we inferred that it likely hydroxylates fraxetin at the structurally analogous position to yield a novel coumarin we have termed sideretin (5-hydroxyfraxetin). We synthesized sideretin from fraxetin using a phthaloyl peroxide-mediated hydroxylation strategy^26^ (Supplementary Fig. 7), and confirmed its identity to **2** (Supplementary Fig. 5b). Sideretin readily oxidizes (e.g. on exposure to air or during column chromatography) to yield **1**, corresponding to the quinone analog (Supplementary Fig. 8), and can be re-reduced back to the hydroquinone upon treatment with H_2_ over Pd.

To complete the biosynthetic pathway of sideretin, we sought an enzyme with scopoletin 8-hydroxylase (S8H) activity capable of producing fraxetin. By searching publicly available transcriptome data, we identified At3g12900, a gene encoding a 2-ODD, which is coexpressed with *CYP82C4*, upregulated under iron deficiency (Supplementary Fig. 3), and has 48% sequence identity to F6’H1 at the amino acid level. We therefore obtained two independent T-DNA insertion lines for functional analysis. Similarly to *CYP82C4*, qPCR analysis confirmed that At3g12900 expression is strongly induced in iron deficient-roots, but abolished in the two selected T-DNA lines (Supplementary Fig. 4b). Consistent with the proposed activity of this enzyme, we observed exudation of scopoletin (**4**) but no further oxidized coumarins in these lines (Fig. 1a). We attempted to express and purify At3g12900 protein in *E. coli* and *N. benthamiana* for *in vitro* biochemical characterization, but could not readily obtain active protein in either case, evidently due to instability (Supplementary Fig. 9).

In lieu of complete *in vitro* biochemical characterization, we attempted to reconstitute the proposed pathway *in vivo* by transiently expressing subsets of pathway enzymes in *N. benthamiana* and analyzing metabolite extracts by LC-MS (Fig. 1b). None of the pathway coumarins (scopoletin, fraxetin or sideretin) are present at detectable levels in *N. benthamiana* leaves expressing only the empty vector. However, the expression of F6’H1 results in the accumulation of scopoletin (Fig. 1b). Notably, leaves expressing both F6’H1 and At3g12900 proteins accumulate fraxetin but not scopoletin, further confirming that At3g12900 functions as a scopoletin 8-hydroxylase (S8H). Finally, expression of all three pathway enzymes leads to accumulation of sideretin but none of the pathway intermediates, further confirming the activity of CYP82C4 (Fig. 1b). Together, these data outline the complete biosynthetic pathway of sideretin from feruloyl-CoA, a lignin precursor found ubiquitously in higher plants (Fig. 1c). Interestingly, expression of F6’H1 also leads to accumulation of esculetin, whose levels are not significantly affected by expression of downstream sideretin pathway enzymes (Fig. 1b). It is not clear whether esculetin arises here via demethylation of scopoletin or side activity of F6’H1 on caffeoyl-CoA, although this latter activity was not seen *in vitro*^21^.

Finally, we profiled the *pdr9*-*2* mutant, which has previously been shown to be defective in export of various coumarins from the roots^18^. The level of sideretin in spent medium for this mutant is approximately 10-fold lower than for wild-type plants (Fig. 1a and Supplementary Fig. 1a), while levels of sideretin glucosides in the roots are significantly elevated (Supplementary Figs. 1b and 2), indicating that PDR9 is the principal sideretin root exporter.

### Chemical characterization of catecholic coumarins

The structure of sideretin is immediately suggestive of its function: catecholic compounds bind to the Fe^3+^ ion, usually forming intensely colored complexes (Supplementary Fig. 10), can reduce Fe(III) to Fe(II), and are therefore well-suited for facilitating iron uptake in Strategy I plants. The possibility that catecholic coumarins are important for iron uptake in *A. thaliana* has been suggested previously, and it is known that *f6’h1* knockouts incapable of producing scopoletin (and therefore, as we have now shown, fraxetin and sideretin) are notably impaired in their iron uptake capacity at elevated pH^19,20^. We therefore hypothesized that sideretin is the major catecholic coumarin facilitating iron uptake from soil in *A. thaliana*. To explore this possibility, we performed a variety of chemical and biological assays with relevant coumarins and pathway mutants.

First, we compared the iron mobilization characteristics of coumarins *in vitro* (Fig. 2). Fraxetin and sideretin, but not scopoletin, can efficiently solubilize iron from hydroxide precipitates (Fig. 2a, b). A kinetic Fe(III) reduction assay further demonstrated that the precise configuration of the catechol moiety and presence of other substituents on the coumarin scaffold have significant effects on Fe(III) reduction kinetics (Fig. 2c). In particular, fraxetin can reduce Fe(III) faster than esculetin and daphnetin (7,8-dihydroxycoumarin), both of which are catecholic coumarins but lack a methoxy substituent. Sideretin, in turn, shows even faster initial kinetics, but reduces less than the expected two equivalents of Fe(III) under the conditions of our assay, possibly due to instability at higher pH (Fig. 2c). We also measured the redox potentials of fraxetin and sideretin by cyclic voltammetry (Fig. 2d and Supplementary Fig. 11): the additional hydroxylation on sideretin lowers the reduction potential by 300 mV with respect to fraxetin (Fig. 2d and Supplementary Fig. 11), consistent with the observation that sideretin is readily oxidized on exposure to air whereas fraxetin is not (Supplementary Fig. 8). In comparison to fraxetin and scopoletin, sideretin also showed high sensitivity to light, reinforcing its high reactivity but weak stability (Supplementary Fig. 12). Together, these data show that catecholic coumarins can both solubilize and reduce Fe(III) and, moreover, that sideretin may be a kinetically and thermodynamically superior reducing agent to fraxetin under certain conditions, potentially explaining the “role” of the additional hydroxylation installed by CYP82C4.

**Figure 2.**
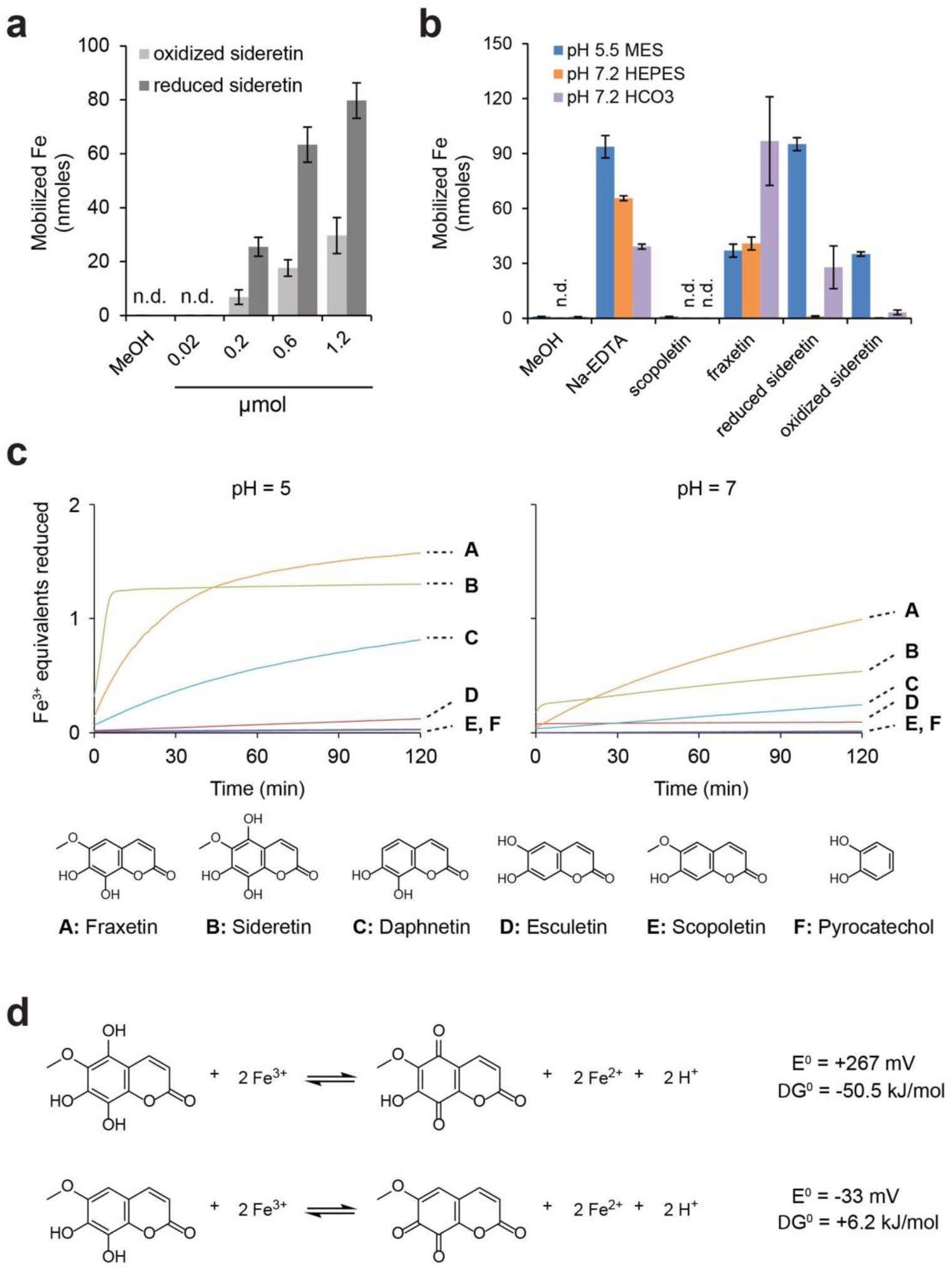
Chemical characterization of the iron mobilization capacity of coumarins. (a, b) Fe(III) mobilization from iron hydroxide precipitates (pH 7.2) by reduced or air-oxidized sideretin at various concentrations (a) and comparison to other catecholic coumarins at 0.6 micromoles (b). Data represent the mean ± s.e. of four replicates (two independent tests). (c) Traces showing Fe(III) reduction kinetics by various compounds at pH 5 or 7, monitored spectrophotometrically by formation of Fe(II)-ferrozine complex. (d) Standard free energy changes and redox potentials for Fe(III) reduction by sideretin and fraxetin, calculated from cyclic voltammetry measurements (see Supplementary Fig. 11) and the literature value of +0.77 V (vs. SHE) for Fe(III) reduction (Fe^3+^ + e^-^ → Fe^2+^).

### Biological characterization of sideretin pathway mutants

In order to test the importance of the sideretin pathway for iron mobilization *in vivo*, we first investigated the localization of *S8H* and *CYP82C4* expression. Similar to *F6’H1*,^19^ the Fe deficiency-dependent upregulation of *S8H* and *CYP82C4* was mainly confined to roots, where the promoter activity of either gene was confined to the primary root and lateral roots, except for the apical root zone (Supplementary Figure 13a, c). At the cell type-specific level, both *S8H* and *CYP82C4* were expressed most strongly in the root epidermal layer under iron-deficient conditions (Supplementary Fig. 13b, d), which is consistent with their role in the biosynthesis of a compound secreted into the rhizosphere. Notably, *F6’H1* is also expressed in the cortex^19^, suggesting that the further conversion of F6’H1-generated scopoletin into fraxetin and sideretin takes place mainly in epidermal cells.

Next, we compared the growth of WT plants with that of various pathway mutants, as well as several other lines known to be impaired in aspects of iron uptake. Due to the photosensitivity of oxidized coumarins, especially sideretin (Supplementary Fig. 12), we initially performed our phenotypic characterization in peat-based soil substrate. In unmodified soil (pH = 5.5), sideretin pathway mutants (*f6’h1*, *s8h*, *cyp82C4*) and the exporter mutant *pdr9*-*2* showed no significant differences from wild-type seedlings in appearance (Fig. 3a), chlorophyll concentration (Fig. 3b), or biomass (Fig. 3c). In contrast, when these plants were grown in soil in which iron availability was decreased by adjusting the pH to 7.2 with calcium carbonate and bicarbonate, *f6’h1*-*1*, *pdr9*-*2* and, crucially, *s8h* lines were visibly more chlorotic and stunted (Fig. 3a), with significantly lower chlorophyll concentrations (Fig. 3b) and biomass (Fig. 3c). Surprisingly, growth of *cyp82C4* lines was not impaired under high pH conditions, and in fact appeared even more robust than that of the wild type, at least in terms of biomass (Fig. 3c). Similar results were obtained on agar plate experiments with FeCl_3_ as the sole iron source, except that the *f6’h1*-*1* line showed mild chlorosis even at pH = 5.5 (Supplementary Fig. 14). To rule out the possibility that exudation of scopoletin alone is capable of mediating iron uptake we tested whether we could complement the phenotypes of *f6’h1* and *s8h* seedlings grown on agar by exogenous addition of various coumarins. We found that fraxetin and sideretin can rescue chlorosis in both *f6’h1*-*1* and *s8h* seedlings, while scopoletin can rescue *f6’h1*-*1* but not *s8h* seedlings (Fig. 3d, e). Thus, whereas externally supplied scopoletin can be further processed to catecholic coumarins in *f6’h1*-*1*, this cannot occur in *s8h* lines, and scopoletin itself clearly cannot reverse the phenotype, even when provided in abundance. Together, these results provide direct evidence for the significance of catecholic compound secretion into the rhizosphere in iron mobilization.

**Figure 3.**
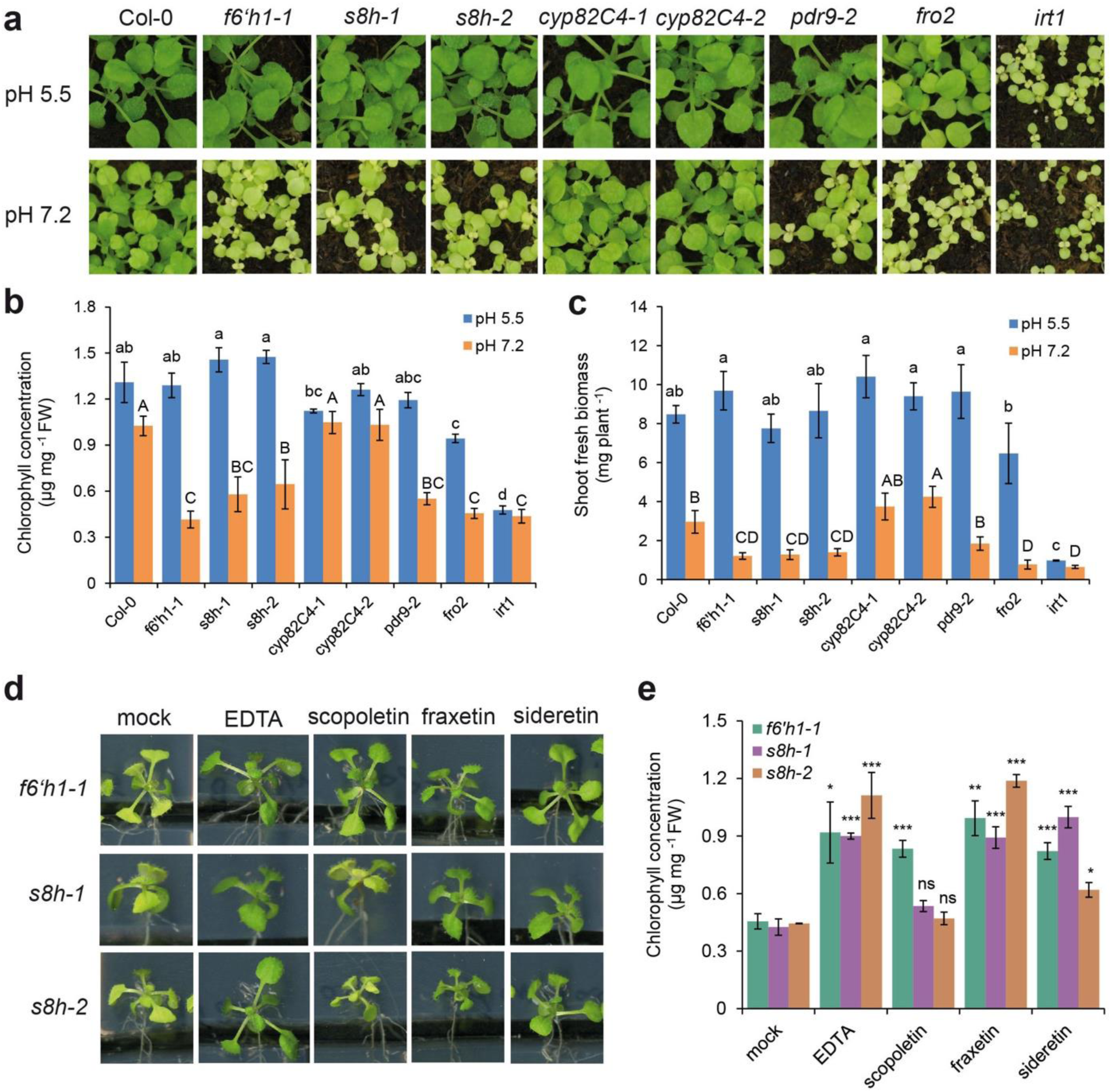
Phenotypic characterization and complementation assays for *s8h*, *cyp82C4* and other mutants under conditions of low Fe availability. (a-c) Appearance of shoots (a), leaf chlorophyll concentration (b) and biomass (c) of WT (Col-0) and various mutant plants grown for 21 days on standard substrate at pH 5.5 or limed substrate at pH 7.2. Data represent mean ± s.d. of 6 biological replicates. Different letters indicate significant differences according to Tukey’s test (p < 0.05). (d, e) Appearance of shoots (d) and leaf chlorophyll concentration (e) of *f6’h1*-*1* and *s8h* mutant lines grown for 7 days on low Fe availability and supplied with indicated compound (150 μM). Data represent mean ± s.e. for 4 biological replicates. ^∗^*p* < 0.05, ^∗∗^*p* < 0.01, ^∗∗∗^*p* < 0.001 and n.s., not significant, two-tailed *t*-test (pairwise comparison with mock treatment for each line separately).

Interestingly, our data seem to suggest that CYP82C4-mediated conversion of fraxetin to sideretin may actually be maladaptive for iron uptake at high pH: sideretin is likely a poorer iron mobilization agent than fraxetin at higher pH (Fig. 2), and may not form stable complexes with Fe(III) (Supplementary Fig. 10). Moreover, *cyp82C4* knockout lines outperform WT plants in certain respects when grown in calcareous soil (Fig. 3c and Supplementary Fig. 14b and c). Given this, we decided to compare the exuded coumarin profile of iron-deficient plants grown at pH 5.7 and pH 7.3 in our hydroponic setup. We found that, at pH = 7.3, while sideretin is still the major exuded coumarin in Col-0 plants, significant amounts of fraxetin, esculetin, and cleomiscosins also accumulate. This contrasts strongly with the exudates collected at pH = 5.7, in which only negligible amounts of coumarins besides sideretin could be detected (Supplementary Fig. 15). This observation suggests that the composition of root exudates of *A. thaliana* responds not only to the plant’s iron nutritional status, but also to soil pH.

### Phylogenetic distribution of sideretin biosynthesis

Finally, we wished to explore whether sideretin biosynthesis is confined to *A. thaliana* and close relatives, or more widespread. To determine the phylogenetic distribution of the sideretin pathway, we used two strategies to identify putative orthologs of the three *A. thaliana* sideretin biosynthetic genes in 54 sequenced plant species. For members of the Brassicaceae family, we performed shared synteny analysis using Genome Browser^27^ (Supplementary Fig. 16), whereas for more divergent species, we performed a reciprocal best BLAST hit search via the KEGG database^28^ (Supplementary File 1). In general, sideretin pathway orthologs appear to be widespread in eudicots, with sporadic loss of all or part of the pathway in certain lineages (Fig. 4a). Interestingly, while *F6’H1* and *S8H* are conserved in all Brassicaceae species analyzed, *CYP82C4* has been independently lost in several members of this family. Furthermore, the basal angiosperm *Amborella trichopoda* contains a *CYP82C4* ortholog, as well as a single *F6’H1/S8H*-like ortholog, However, all pathway orthologs are conspicuously absent from grassy (Strategy II) plants which suggests Fe acquisition via coumarins and phytosiderophores do not likely co-exist in nature. Given these facts, coupled with the high sequence similarity of F6’H1 and S8H, we speculate that this pathway arose early in angiosperm evolution, partly via a gene duplication event that gave rise to the *F6’H1/S8H* paralog pair, but may have been supplanted by alternative iron mobilization strategies in various lineages.

**Figure 4.**
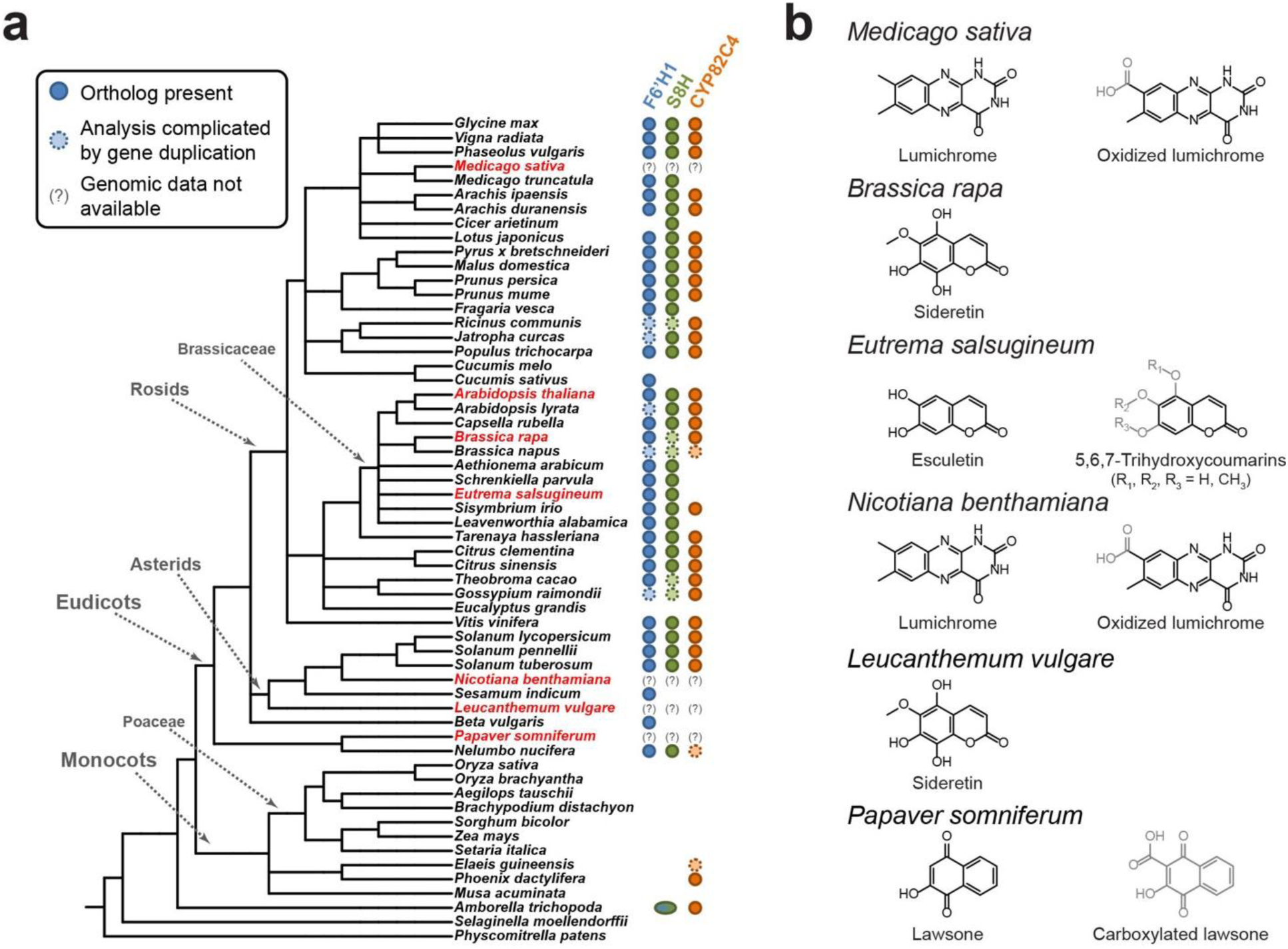
Phylogenetic distribution of sideretin pathway enzyme orthologs and Fe deficiency-induced small molecule exudation in various plants. (a) Presence of sideretin pathway orthologs in sequenced plants, inferred from shared synteny or reciprocal best BLAST hit analysis. *Amborella trichopoda*, the basal angiosperm, contains a single *f6’h1/s8h*-like ortholog. Species highlighted in red were those profiled for small molecule exudation under iron deficiency in this study. (b) Structures of major redox-active and/or iron-binding compounds secreted in response to iron deficiency in profiled species.

To experimentally validate some of the conclusions drawn from the phylogenetic analysis, we profiled the root exudates of several plant species amenable to axenic growth in suitably modified versions of our hydroponic platform under iron-abundant and iron-depleted conditions at pH = 5.7. We first investigated whether loss of *CYP82C4* in a member of the Brassicaceae family correlates with loss of sideretin production. Towards this end, we grew *Brassica rapa* and *Eutrema salsugineum*, which have, respectively, retained and lost *CYP82C4*. Notably, *E. salsugineum* is an extremophile that preferentially inhabits alkaline salt flats^29^, unlike *A. thaliana* or *B. rapa*. As expected, *B. rapa* exudes sideretin under iron-depleted conditions (Supplementary Fig. 17a and Fig. 4b), whereas *E. salsugineum* exudes a complex mixture of oxidized coumarins, primarily esculetin, isoscopoletin, and various methylated isomers of 5,6,7-trihydroxycoumarin (Supplementary Fig. 17b and Fig. 4b), but no further hydroxylated coumarins like sideretin. At present, how these coumarins are biosynthesized in *E. salsugineum* is unclear. However, this finding does further suggest that different chemical strategies – in this case, exudation of oxidized coumarins with different iron binding and/or redox properties – likely evolved to fit different species’ ecological requirements.

We further profiled four different eudicot species, *Medicago sativa*, *Nicotiana benthamiana*, *Leucanthemum vulgare*, and *Papaver somniferum* for small molecule exudation in response to iron deficiency. We found that *L. vulgare* exudes sideretin (Supplementary Fig. 18a and Fig. 4b), demonstrating that the complete biosynthetic pathway was in place before the divergence of the rosids and asterids approximately 125 million years ago^30^. In contrast, both *M. sativa* and *N. benthamiana* exude lumichrome, the redox-active moiety of the flavin cofactors, as well as a further oxidized derivative whose structure is not entirely clear (Supplementary Fig. 18b, c and Fig. 4b). The exudation of modified riboflavins in response to iron deficiency has previously been demonstrated in *Beta vulgaris*^31^, which lacks S8H and CYP82C4 orthologs, and has recently been implied by transcriptomics analysis in *Medicago truncatula*^15^. Intriguingly, we also observed iron deficiency-induced exudation of acetosyringone by *N. benthamiana* and flavonoids by *M. sativa* (Supplementary Fig. 18b and c). These compounds do not possess obvious redox-active or iron-binding moieties, but are known to mediate plant-microbe interactions^32^. Finally, we detected lawsone in *P. somniferum* exudates (Supplementary Fig. 18d and Fig. 4b), as well as a carboxylated derivative, which we suspect is produced via hydroxylation of 1,4-dihydroxy-2-naphthoic acid, a phylloquinone precursor that is ubiquitous in plants. Remarkably, lawsone possesses the same 1,2,4-trihydroxybenzene substitution pattern seen in sideretin, but on a naphthalene rather than a coumarin scaffold. Although we have not yet performed detailed biochemical characterization of these compounds, the fact that all eudicot species tested by us so far exude redox-active and/or iron-binding molecules under iron deficiency suggests this phenomenon may be a general and previously underappreciated feature of Strategy I iron uptake.

## Discussion

While root exudation of compounds capable of binding and/or reducing iron in various Strategy I plants was first observed over four decades ago^33,34^, the relative importance of this phenomenon in iron mobilization and the underlying biochemical mechanisms have long remained obscure. More recently, studies with *A. thaliana* have shown that synthesis^19,20^ and export^18,35^ of coumarins are crucial for efficient iron uptake under iron-limiting conditions, but still failed to explain the mechanisms underlying this effect. Here, we have identified a widely conserved pathway for the biosynthesis of catecholic coumarins in Strategy I plants and demonstrated the biochemical basis for the role of these compounds in iron mobilization from soil.

The secretion of redox-active metabolites from plant roots is reminiscent of the small molecule electron shuttles used by certain bacterial species for mineral reduction. For instance, phenazines produced by *Pseudomonas* are thought to contribute to iron acquisition, while flavins serve as electron shuttles for anaerobic *Shewanella*, enhancing the rate of extracellular respiration of Fe(III)^36,37^. Although the reactivity of sideretin to light and oxygen has made it difficult to probe whether a related mechanism of redox cycling occurs near plant roots, our findings are consistent with a role for sideretin in extracellular electron transport. It will be interesting to probe whether plant-derived catecholic coumarins act alone, or instead link root surface reductases to otherwise inaccessible pools of precipitated Fe(III).

While our work paves the way for a deeper mechanistic understanding of the role of small molecules in iron mobilization, several unresolved questions remain. In particular, the importance of CYP82C4-mediated conversion of fraxetin to sideretin is unclear, and we have thus far been unable to find any beneficial phenotype for WT plants in comparison to *cyp82C4* knockouts. We suspect that the unique redox capacity of sideretin and the marked preference for its production under low pH conditions are relevant under certain natural conditions, which however may be difficult to capture experimentally. It is possible that at high pH, production of less oxidized coumarins like fraxetin and esculetin is preferred simply due to their enhanced stability and ability to stably chelate Fe(III), but the relative importance of chelation and reduction by catecholic compounds may be difficult to disentangle *in vivo*.

Further, despite the seemingly wide conservation of the sideretin pathway in eudicots, it is apparent that numerous additional chemical strategies for iron mobilization exist, and a broader search than the one we have undertaken here will likely reveal others. Future efforts will entail uncovering the biosynthesis of these additional molecules and clarifying the link between specific chemical strategies and iron mobilization capacity under various environmental conditions affecting iron bioavailability.

To date, most efforts in understanding soil iron uptake limitations have focused on the role of soil pH, ignoring many other potentially relevant factors, such as interactions with soil organic matter or other metals like Zn(II) or Mn(II), which are not redox-active in soil but are also imported by IRT1. With a detailed biochemical understanding of the pathways involved, we can now hope to approach some of these problems. Ultimately, we hope that this knowledge can be applied towards engineering crop varieties with enhanced iron acquisition capacity, enabling growth in otherwise unsuitable environments.

## Acknowledgements

This work was supported by the US National Institutes of Health grant DP2 AT008321 and an HHMI-Simons Faculty Scholar Award to (E.S.S.). and by a grant of the Deutsche Forschungsgemeinschaft to N.v.W. (WI1728/21-1). J. R. was supported by an NIH biotechnology training grant (T32 GM008412-20). We thank Nicole Schmid and Mathias Voges for valuable discussions and help with experiments, Sean Elliott for advice on redox potential measurements, Scott Fendorf for helpful discussions on rhizosphere Fe, M. Kevin Brown and Justin Du Bois for suggestions for chemical synthesis of sideretin, and Thomas Veltman for help with CV measurements.

---

**Supplementary Figure 1.**
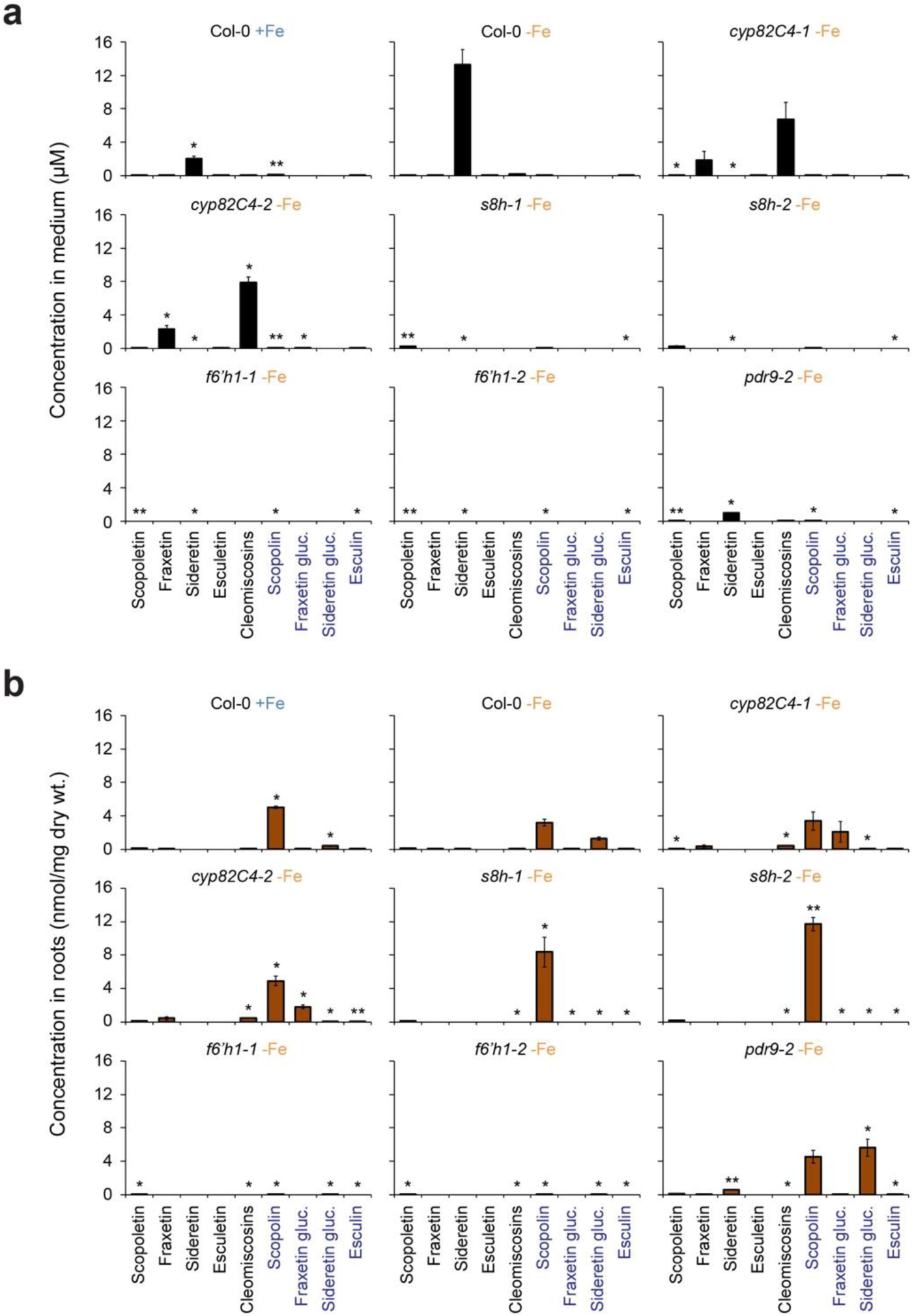
Levels of oxidized coumarins and their glucosides produced by sideretin pathway mutants. (a, b) Compound levels in spent medium (a) and roots (b) of indicated lines after 12 d of hydroponic growth at pH = 5.7. Data shown are mean ± s.d. for three biological replicates. ^∗^*p* < 0.05, ^∗∗^*p* < 0.005, two-tailed *t*-test; all comparisons with respect to amounts in the Col-0-Fe condition. Levels of fraxetin glucosides and sideretin glucosides are approximate: for fraxetin glucosides, the ionizability of all isomers was assumed to be the same as for the commercially available fraxetin-8-*O*-glucoside (fraxin), while for sideretin glucosides, where no standard is available, the ionizability was estimated by comparing glucosidase-untreated and glucosidase-treated samples in a reconstitution experiment in *N. benthamiana*.

**Supplementary Figure 2.**
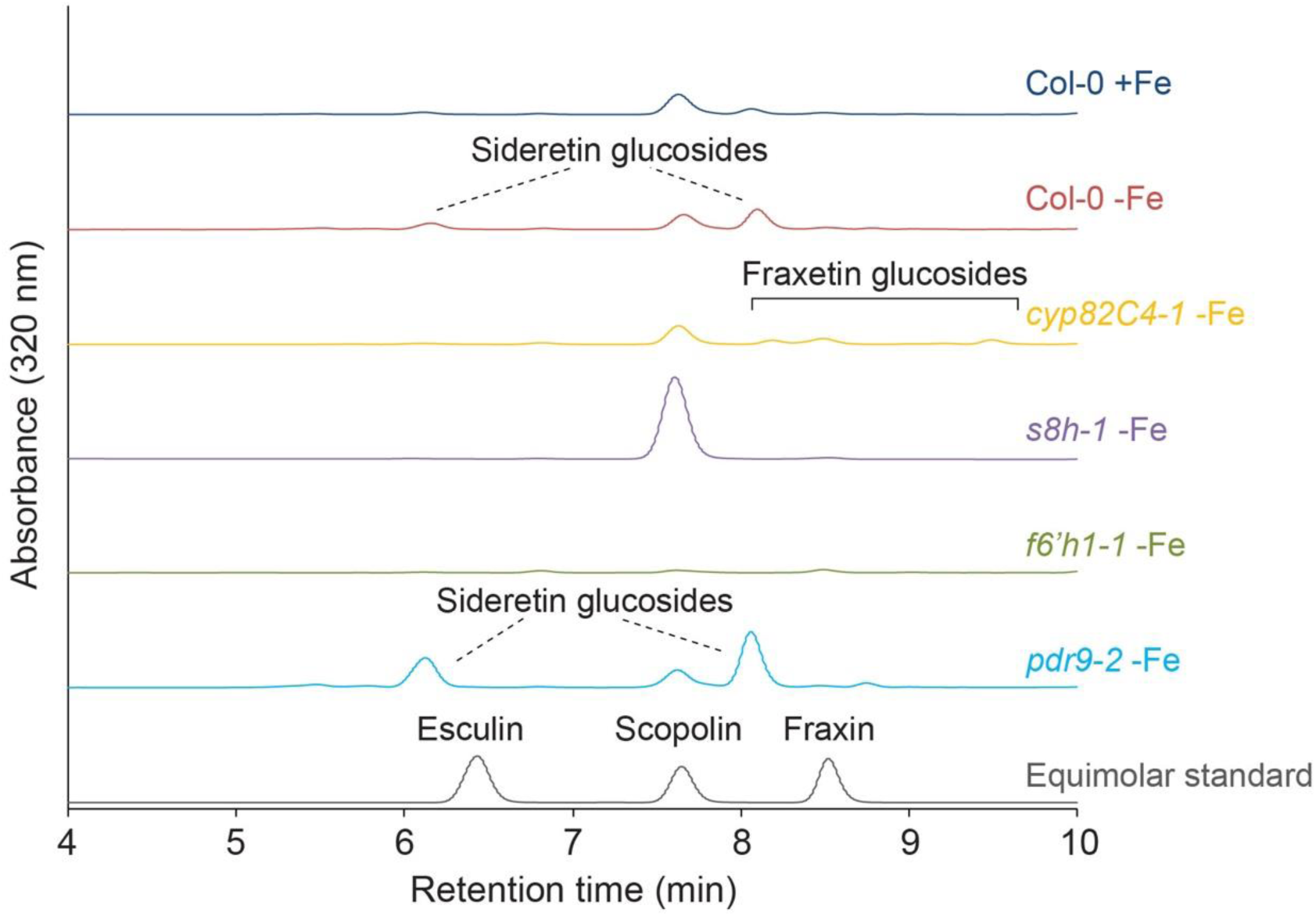
UV-Vis absorbance traces of root extracts for WT and mutant plants. Traces are representative of three biological replicates.

**Supplementary Figure 3.**
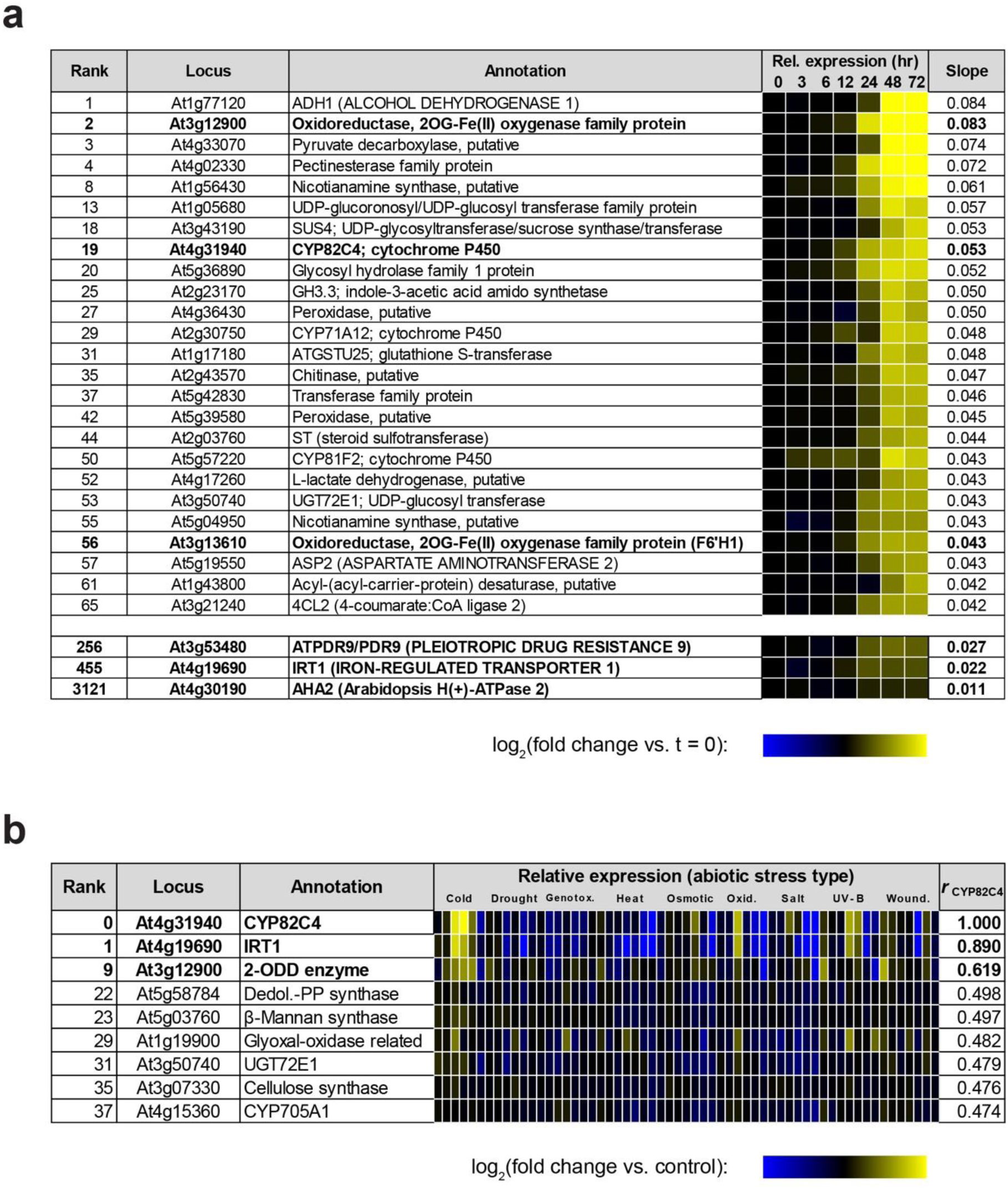
Transcriptomics analysis in *A. thaliana* for identification of candidate sideretin biosynthetic enzymes. (a) Relative induction of gene expression in roots under iron deficiency. Genes are sorted by slope of log-normalized fold change vs. time. (b) Gene expression under various stress conditions in roots. Genes are sorted by correlation (Pearson’s *r*) with *CYP82C4*. In both panels, only enzyme-encoding genes are shown.

**Supplementary Figure 4.**
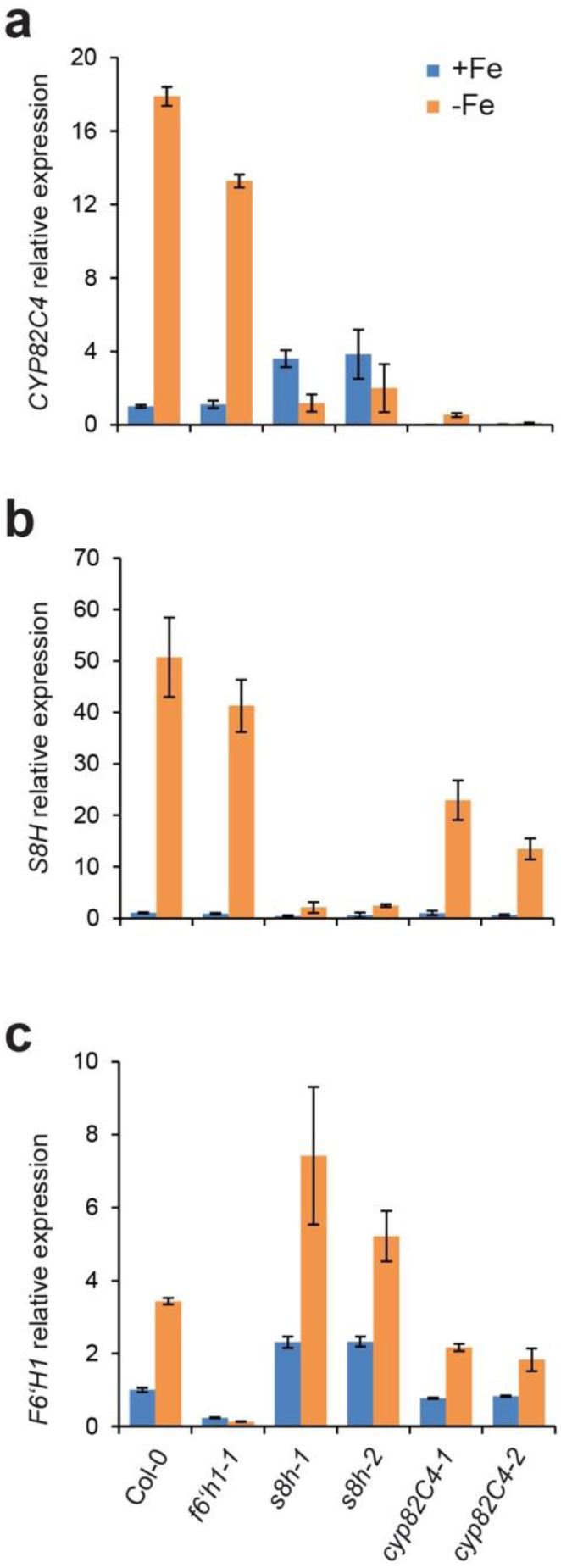
Iron-dependent regulation of *F6‘H1*, *S8H*, and *CYP82C4* expression in roots. (a-c) Relative expression levels analyzed by qPCR of *CYP82C4* (a), *S8H*(At3g12900) (b) and *F6’H1* (c) in roots of wild-type (Col-0) or indicated mutants after six days of treatment. Seedlings were pre-cultured for 7 days on half-strength MS medium (75 μM Fe-EDTA) and transferred to one-half-strength MS medium containing 75 μM Fe-EDTA (+Fe) or no added Fe plus 15 μM of the Fe chelator ferrozine (-Fe). Expression of the housekeeping gene UBQ2 served as reference in all assays. Data represent the mean ± s.e. of four biological replicates.

**Supplementary Figure 5.**
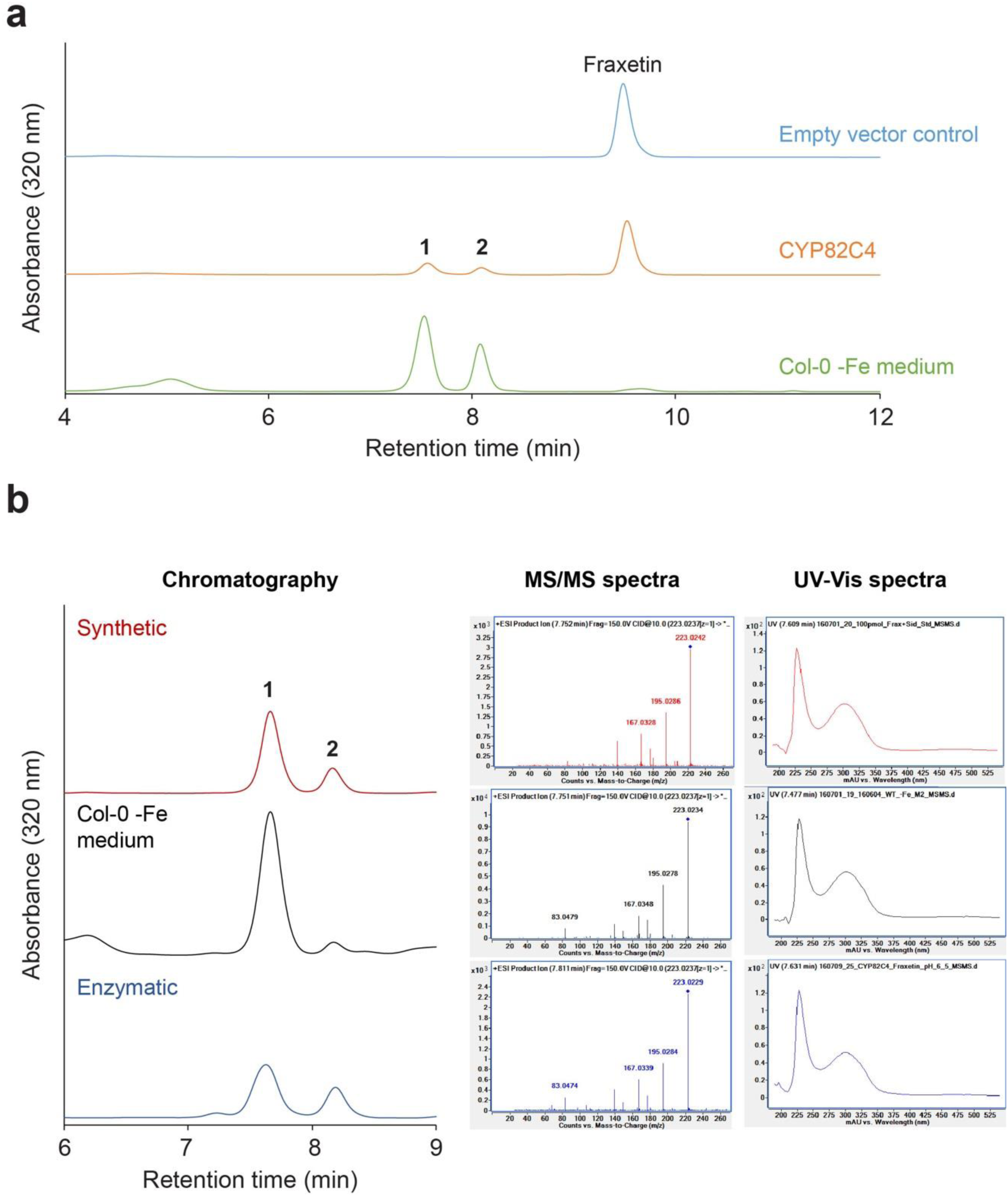
Incubation of fraxetin with CYP82C4 yields sideretin *in vitro*. (a) UV-Vis absorbance traces of enzymatic reactions with fraxetin show that the major peaks seen in WT exudates (**1**, **2**) arise via hydroxylation of fraxetin by CYP82C4. (b) Retention time comparisons, MS/MS fragmentation patterns, and UV-Vis absorbance profiles establish the identity of sideretin from various sources. MS/MS (10 V collision energy) and UV-Vis profiles are for the oxidized species (**1**).

**Supplementary Figure 6.**
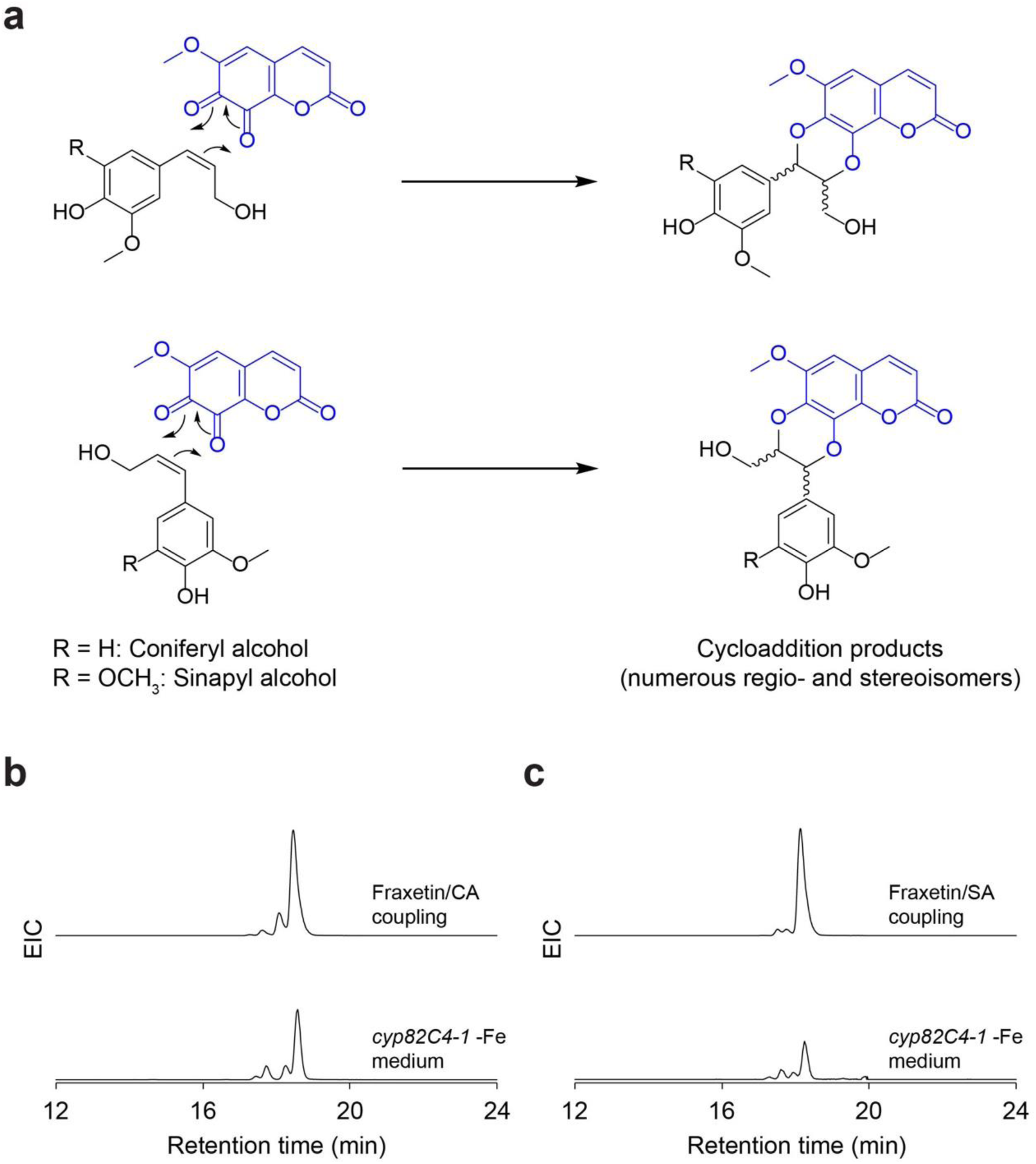
Oxidized fraxetin reacts with monolignols to generate cleomiscosins. (a) Presumed [4+2] cycloaddition reactions of oxidized fraxetin with alkene-containing monolignols. (b, c) Extracted ion chromatograms comparing retention times of compounds observed in *cyp82C4*-*1* spent medium and products of the oxidative coupling of coniferyl alcohol (b) (Extracted m/z [M+H]^+^ = 387.1074; C_20_H_18_O_8_) and sinapyl alcohol (c) (Extracted m/z [M+H]^+^ = 417.1180; C_21_H_20_O_9_). Note that non-specific addition would give rise to four separable isomer peaks (that is, four pairs of enantiomers) for each synthesis, as observed in the EIC traces.

**Supplementary Figure 7.**
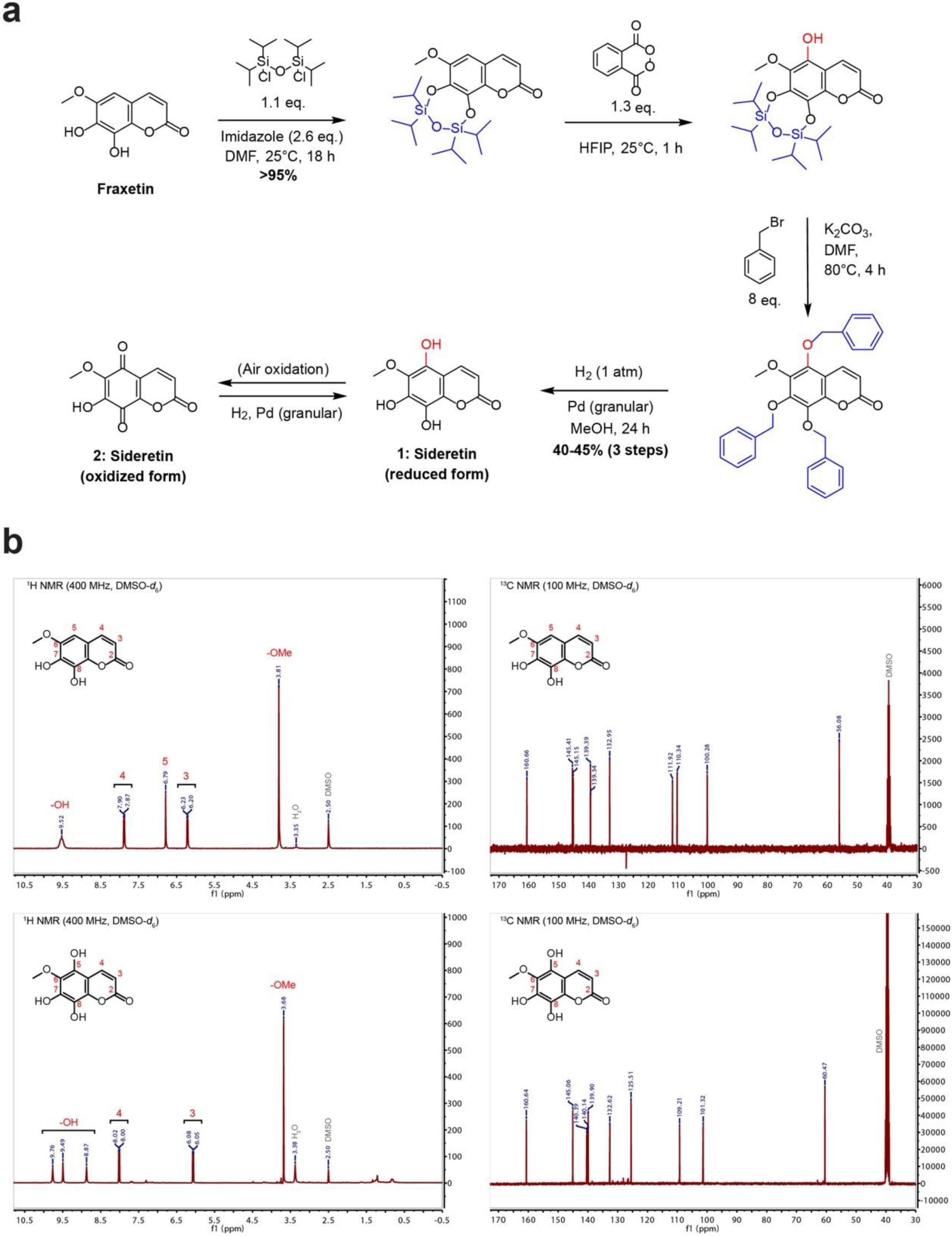
Synthesis of sideretin from fraxetin. (a) Overall synthetic scheme for sideretin via phthaloyl peroxide hydroxylation of fraxetin. (b) Comparison of ^1^H and ^13^C NMR spectra for fraxetin standard and synthetic sideretin. Loss of the unique fraxetin aromatic resonance (δ = 6.79) in the sideretin ^1^H spectrum demonstrates that sideretin is hydroxylated at carbon 5 (standard coumarin carbon atom numbering shown).

**Supplementary Figure 8.**
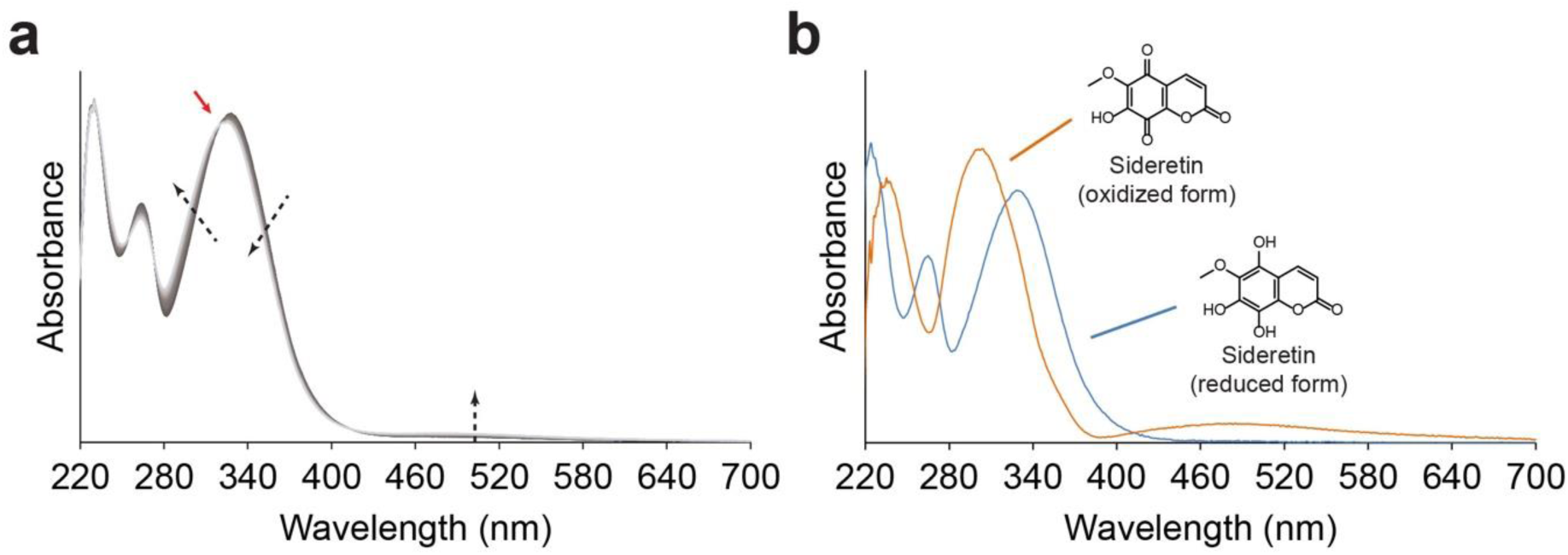
UV absorbance properties of sideretin. (a) A scanning kinetics experiment shows the progress of sideretin oxidation in air. Black arrows indicate change in absorbance with time; the red arrow indicates an isosbestic point at 320 nm. (b) UV absorbance spectra for the quinone and catechol forms of sideretin. The catechol spectrum was obtained for pure compound under anoxic conditions, while the quinone spectrum was found by deconvolution from the scanning kinetics data.

**Supplementary Figure 9.**
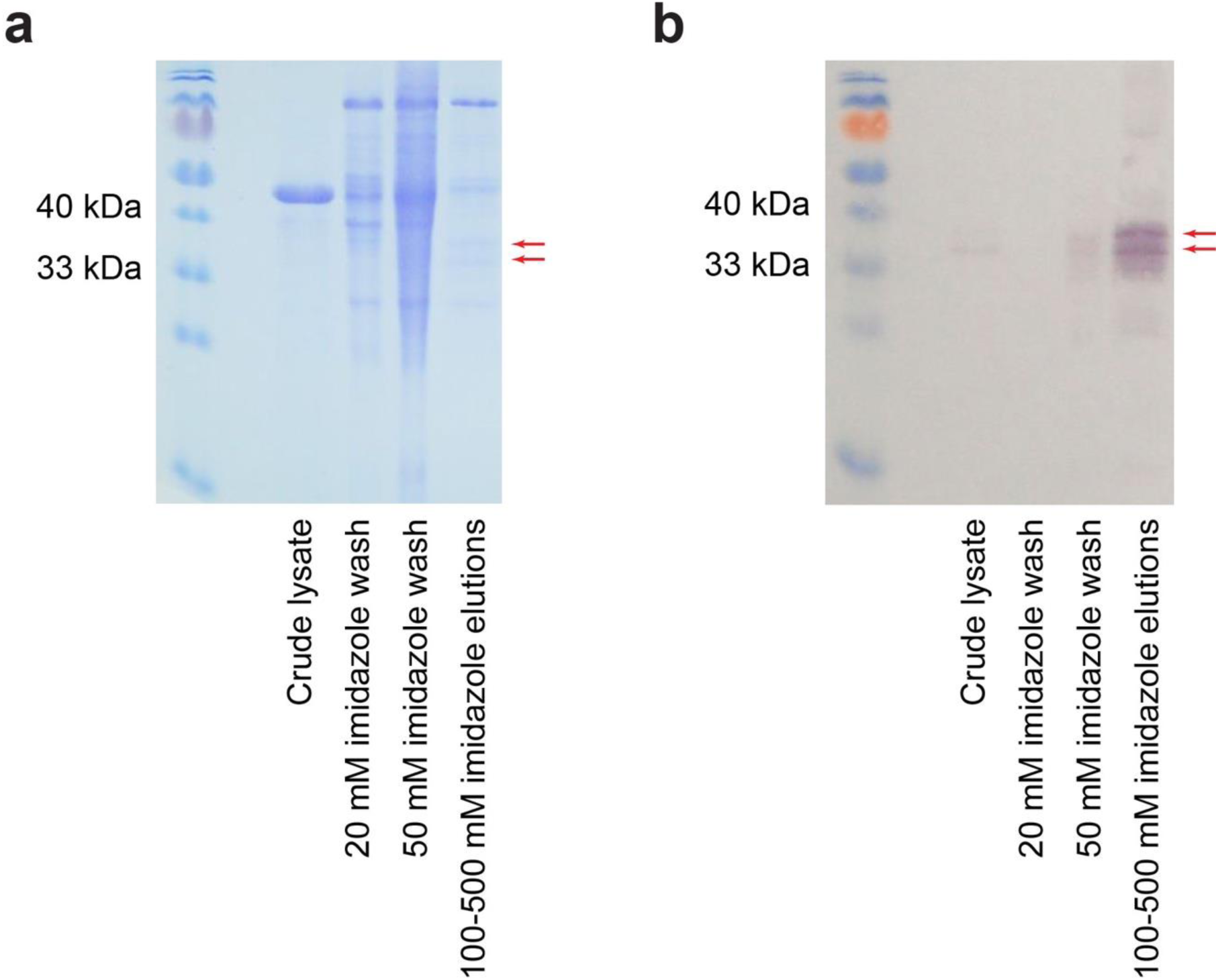
Partial degradation of *A. thaliana* S8H protein heterologously expressed in *N. benthamiana* leaves. (a, b) Coomassie-stained protein gel (a) and anti-6xHis Western blot (b) of S8H protein nickel affinity chromatography purification fractions. The predicted molecular weight of the full-length product is 41 kDa.

**Supplementary Figure 10.**
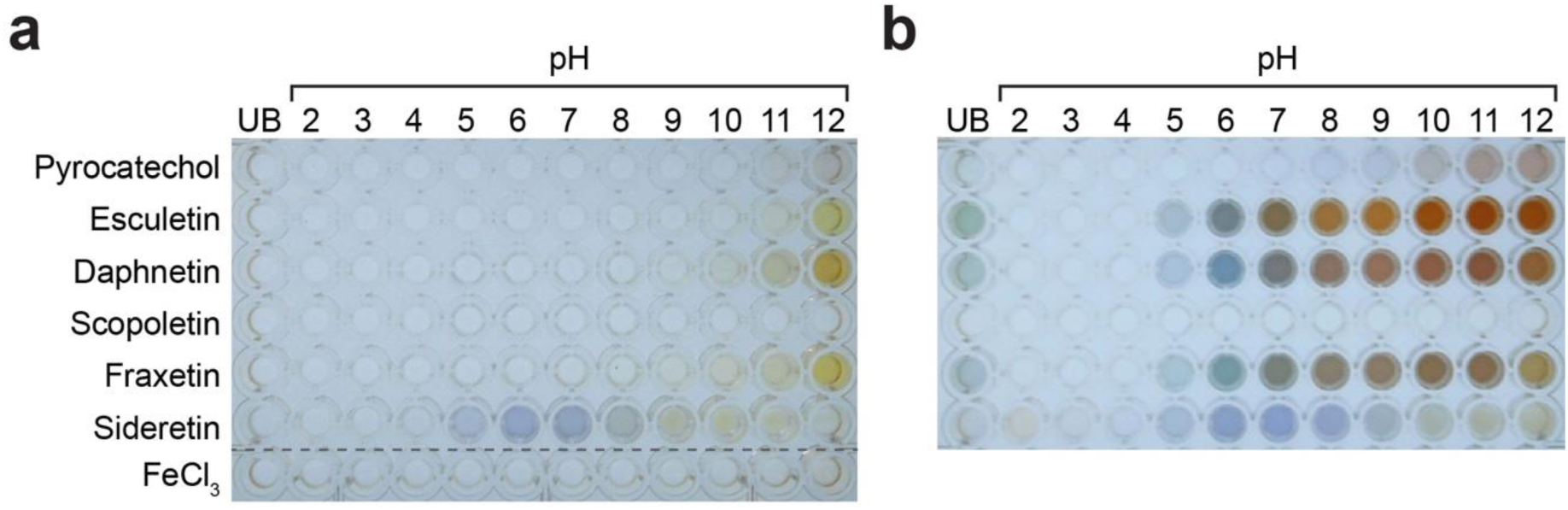
Formation of catecholic coumarin-Fe(III) complexes. (a, b) Appearance of 750 μM compound standards (250 μM for FeCl_3_) in aqueous solution buffered with Britton-Robinson buffer at the indicated pH (a), or 5 min after addition of 250 μM FeCl_3_ (b). Note that scopoletin, which lacks a catechol moiety, does not form a colored complex with Fe(III) at any pH; furthermore, sideretin initially forms colored complexes of comparable intensity to the other catecholic coumarins, but the color rapidly fades, suggesting that sideretin-Fe(III) complexes are unstable. UB: unbuffered.

**Supplementary Figure 11.**
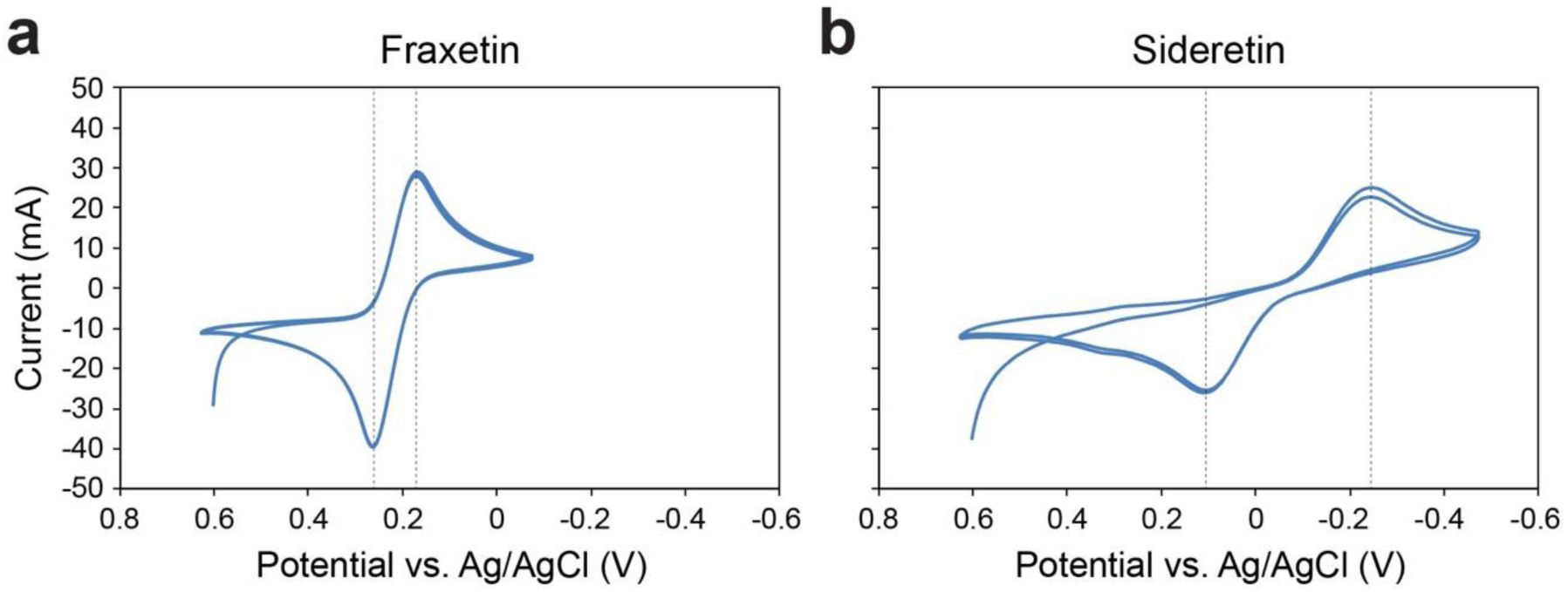
Cyclic voltammetry of fraxetin and sideretin. (a, b) Voltammetry traces for fraxetin (a) and sideretin (b) were obtained in aqueous phosphate buffer at pH 6.5. The average of the two potentials at maximal current (indicated by dashed lines) is, after correction (see Materials and Methods), a good estimate of the standard redox potential. The standard redox potentials (vs. SHE) calculated in this manner are +0.803 V for fraxetin and +0.503 V for sideretin.

**Supplementary Figure 12.**
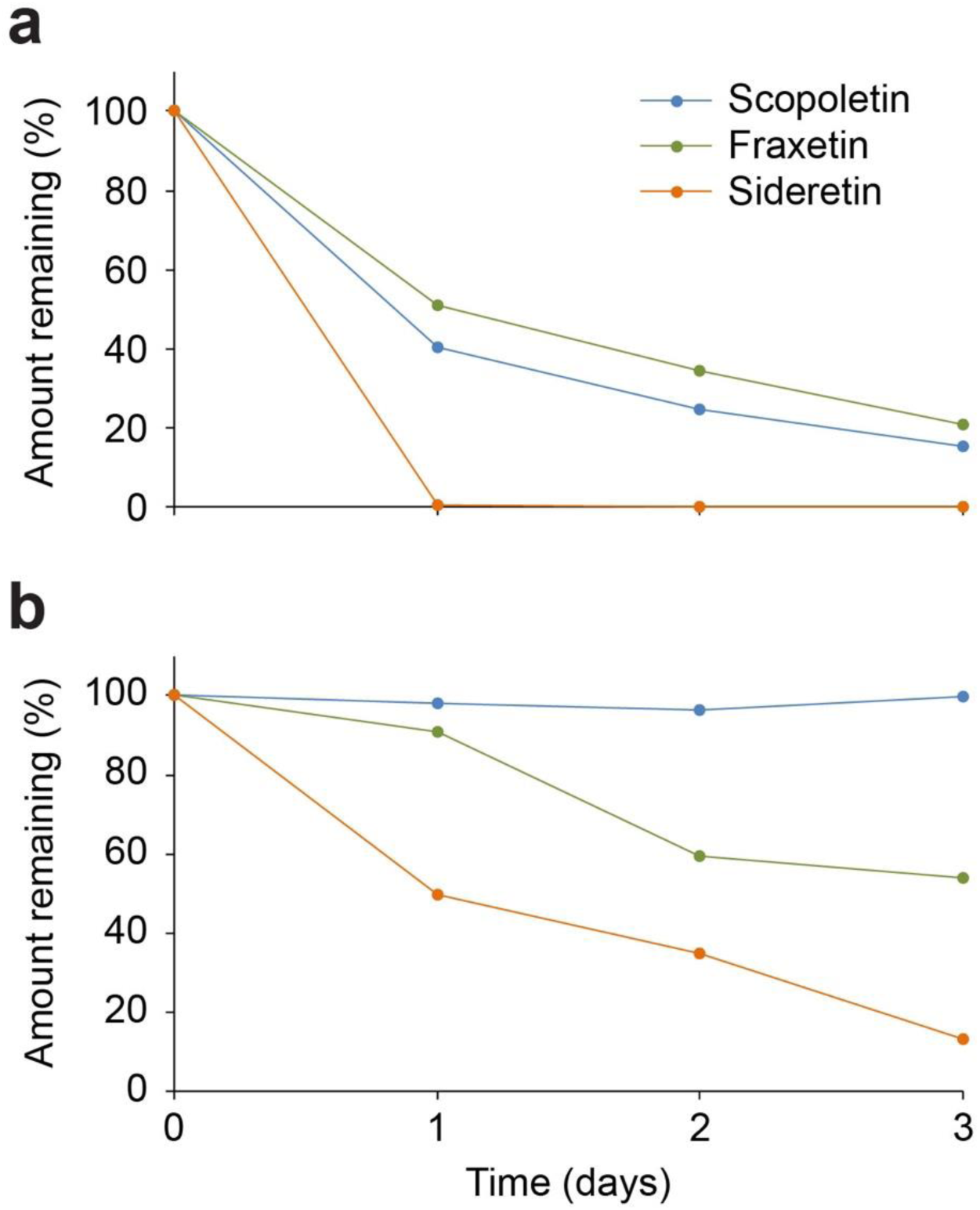
Photosensitivity of coumarins in hydroponic medium. (a, b) Coumarin degradation over time in MS medium, either under exposure to standard light conditions (16 h day cycle) in a growth chamber (a) or no light exposure (b).

**Supplementary Figure 13.**
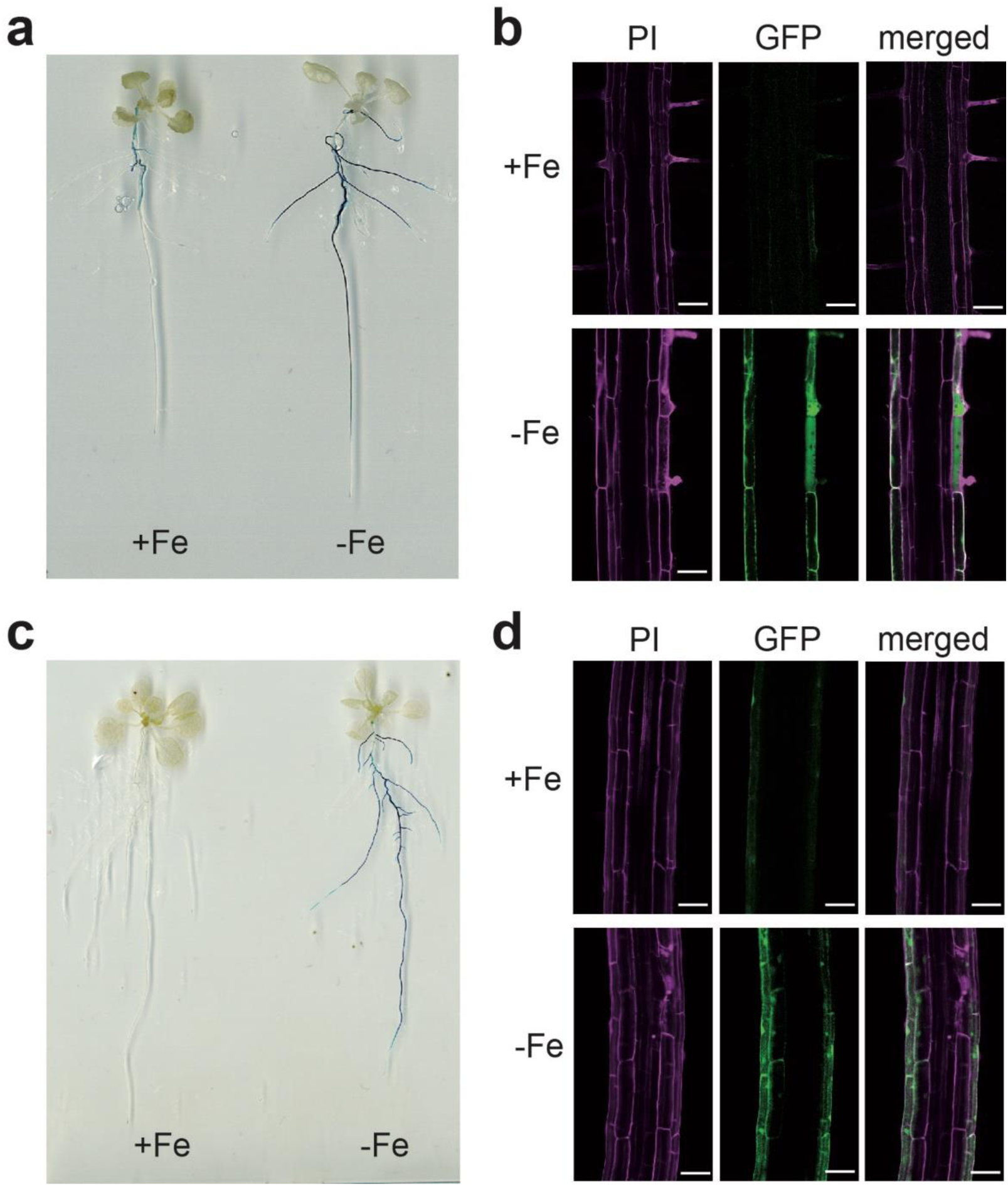
Tissue- and cell-type specific localization of *S8H* and *CYP82C4* expression. (a, b) proS8H-dependent GUS activity (a) and GFP expression (b) in Fe-sufficient or Fe-deficient plants. (c, d) proCYP82C4-dependent GUS activity (c) and GFP expression (d) in Fe-sufficient or -deficient plants. Plants were precultured in half-strength MS medium (75 μM Fe-EDTA) medium for 7 days and then transferred to half-strength MS medium with 75 μM Fe-EDTA (+Fe) or no added Fe plus 15 μM of ferrozine, a strong Fe chelator (-Fe). All plates were buffered at pH = 5.5 with 2.5 mM MES. In b and d, representative root sections from one representative transgenic line (n = 8) are shown. Scale bars = 50 μm. Pink fluorescence derives from propidium iodide (PI) staining of cell walls.

**Supplementary Figure 14.**
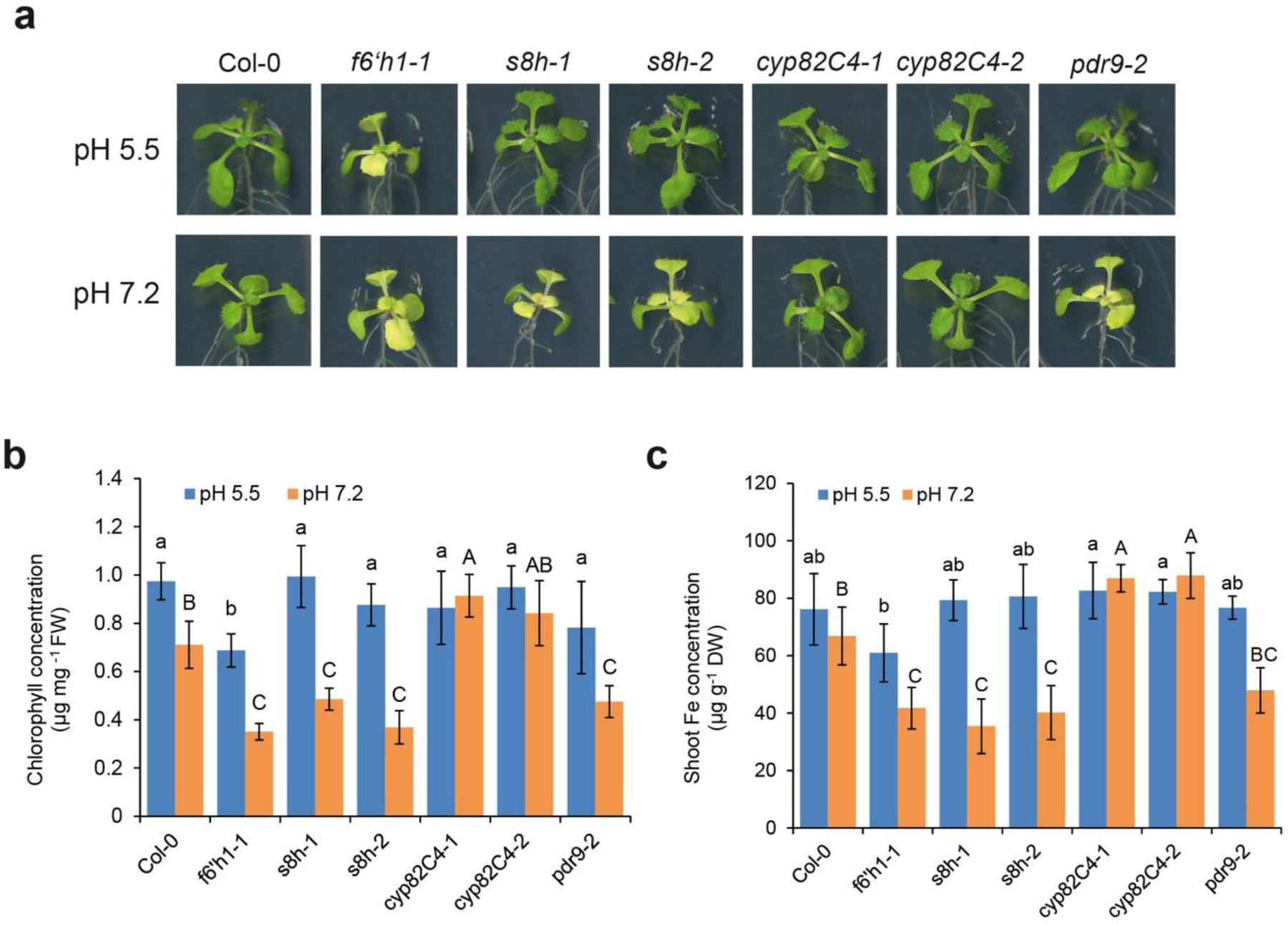
Phenotypic characterization of sideretin pathway mutants on agar. (a-c) Appearance of shoots (a), leaf chlorophyll concentration (b) and shoot Fe concentration (c) of WT (Col-0) and various mutant plants grown for 6 days on low Fe availability at the indicated pH. Results shown represent the mean ± s.d. of four biological replicates (two independent experiments). Different letters indicate significant differences according to Tukey’s test (*p* < 0.05).

**Supplementary Figure 15.**
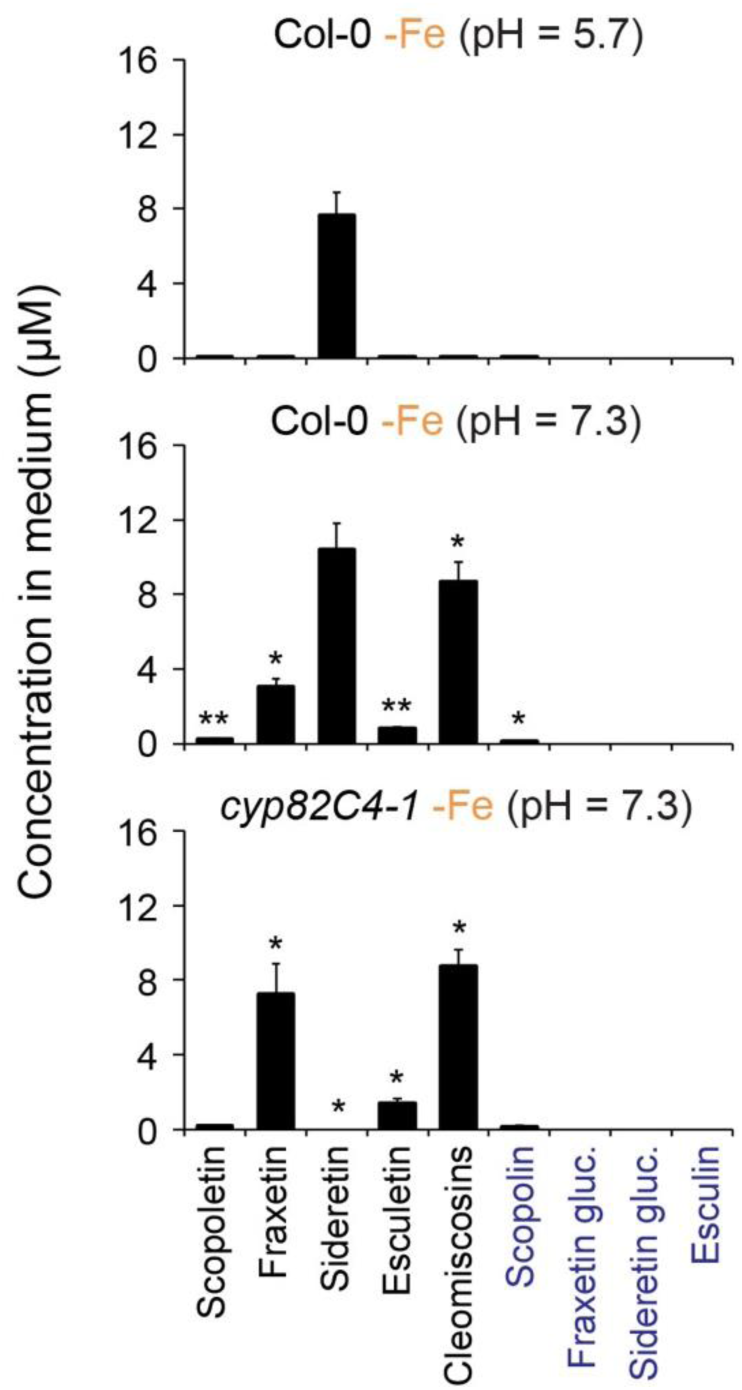
Comparison of root-exuded coumarin levels at pH 5.7 and pH 7.3. Compound levels in spent medium after 12 d of growth at the indicated pH. Data shown are mean ± s.d. for three biological replicates. ^∗^*p* < 0.05, ^∗∗^*p* < 0.005, two-tailed *t*-test; all comparisons with respect to amounts in Col-0 -Fe (pH = 5.7) condition. Levels of compounds for which exact standards are not available were determined as described in Supplementary Fig. 1.

**Supplementary Figure 16.**
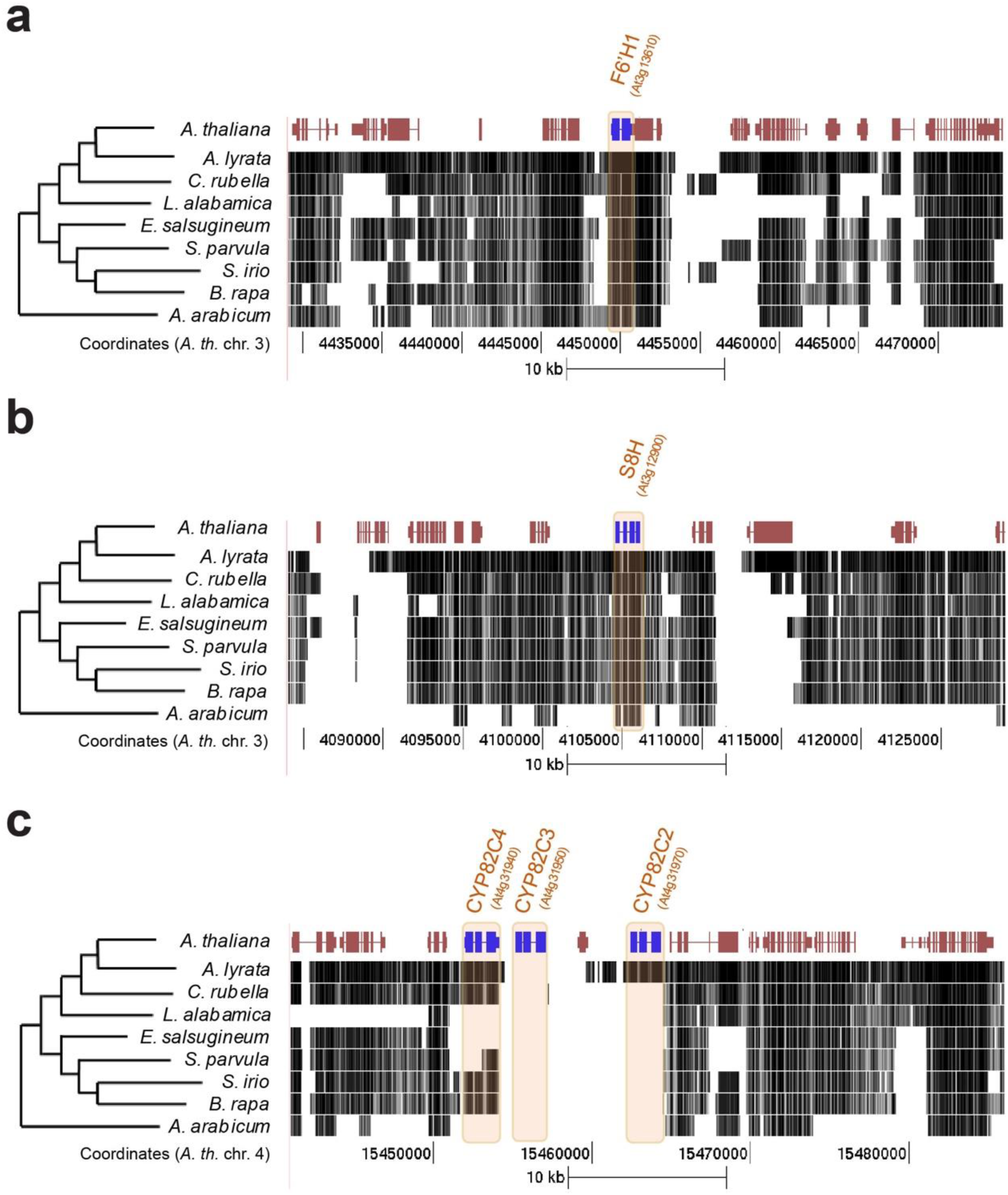
Comparative genomics for sideretin pathway genes in the Brassicaceae family. (a-c) Multiple genome alignments against *A. thaliana* sideretin pathway enzyme genes for a set of 8 sequenced plants. For *A. thaliana*, gene models from the TAIR9 release are shown. Figures were generated using the USCS Genome Browser (CIT). The *F6’H1*(a) and *S8H* (b) genes appear to be well conserved among all species in this alignment, whereas the CYP82C locus (c) shows a complex evolutionary history, marked by independent chromosomal deletions in various lineages, as well as tandem duplication and neofunctionalization in *Arabidopsis* species. Notably, a complete CYP82C4 gene is absent from *L. alabamica*, *E. salsugineum*, *S. parvula*, and *A. arabicum*.

**Supplementary Figure 17.**
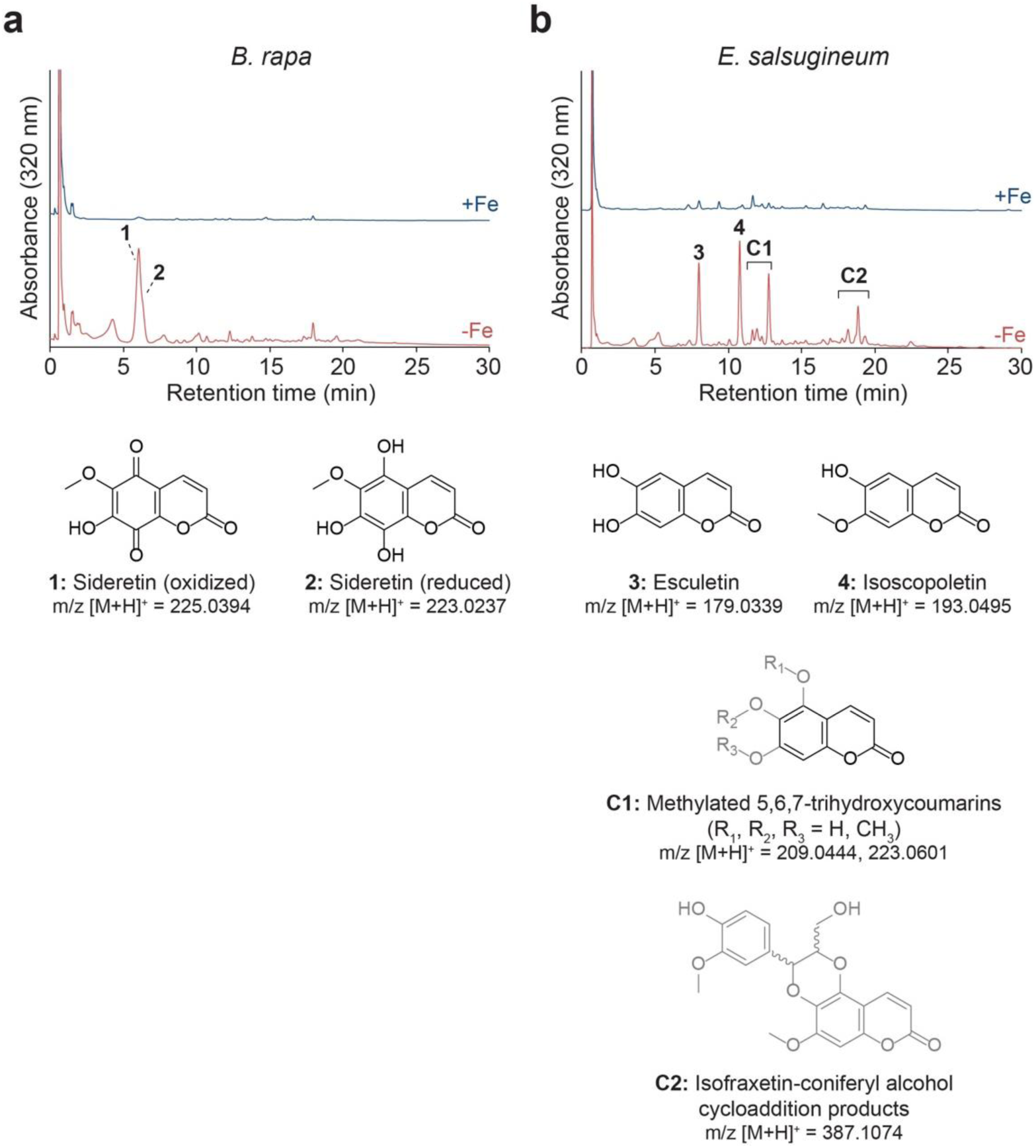
Iron deficiency-induced root exudation of oxidized coumarins by other species in the Brassicaceae family. (a, b) Comparison of UV-Vis absorbance traces for spent medium extracts of hydroponically grown *B. rapa* (a) and *E. salsugineum* (b) grown under Fe-sufficient (+Fe) or Fe-deficient (-Fe) conditions; data are representative of three biological replicates. Structures of compounds shown in black were confirmed by comparison with authentic standards, while those in gray were inferred from m/z values and chromatographic properties. In particular, for *E. salsugineum* (b), peak cluster **C1** includes various methylated isomers of a triply hydroxylated coumarin, but none of these correspond exactly to fraxetin or its methylated derivatives; likewise, cluster **C2** consists of peaks with masses matching those of fraxetin-coniferyl alcohol cycloaddition products (see Supplementary Fig. 6), but distinct by chromatography and MS/MS analysis. In conjunction with the high amount of isoscopoletin observed in these exudates, it is most likely that these peaks correspond to isofraxetin (5,6-dihydroxy-7-methoxycoumarin) and derivatives, for which standards were not easily obtainable.

**Supplementary Figure 18.**
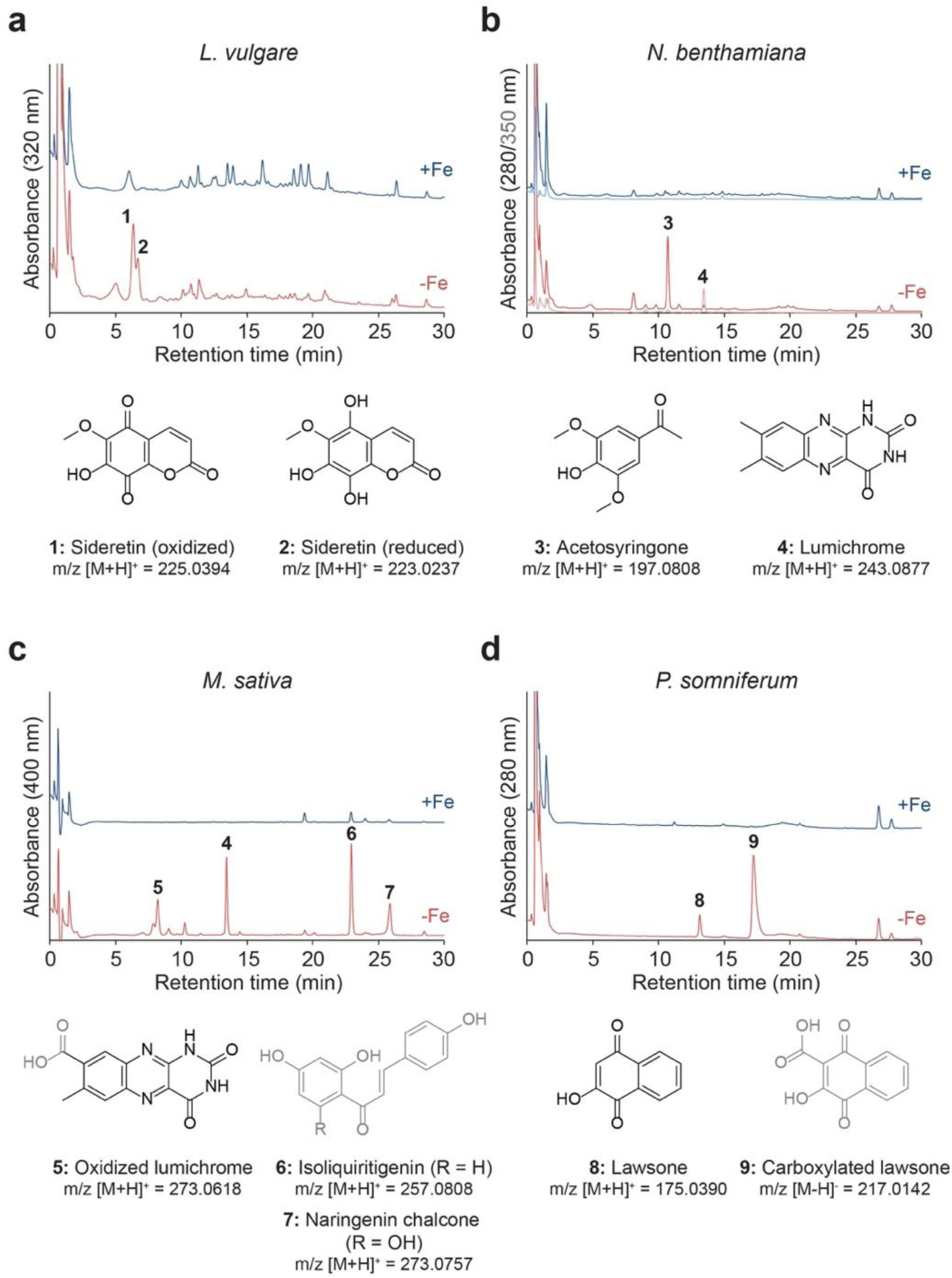
Iron deficiency-induced root exudation of structurally diverse compounds by various eudicot species. (a-d) Comparison of UV-Vis absorbance traces for spent medium extracts of hydroponically grown *L. vulgare* (a), *N. benthamiana* (b), *M. sativa* (c), and *P. somniferum* (d) grown under Fe-sufficient (+Fe) or Fe-deficient (-Fe) conditions; data are representative of three biological replicates. As in Supplementary Fig. 17, structures shown in black were confirmed by comparison with authentic standards, while those in gray were inferred from m/z values and chromatographic properties. For *M. sativa* (c), assignment of the two late-eluting compounds **6** and **7** as isoliquiritigenin and naringenin chalcone, respectively, was additionally motivated by previous reports of their occurrence in this species, as well as distinctive UV-Vis absorbance profiles.

## References

1. Hänsch, R. & Mendel, R. R. Physiological functions of mineral micronutrients (Cu, Zn, Mn, Fe, Ni, Mo, B, Cl). Current Opinion in Plant Biology 12, 259–266 (2009).

2. Balk, J. & Pilon, M. Ancient and essential: the assembly of iron–sulfur clusters in plants. Trends in Plant Science 16, 218–226 (2011).

3. Palmer, C. M. & Guerinot, M. L. Facing the challenges of Cu, Fe and Zn homeostasis in plants. Nat Chem Biol 5, 333–340 (2009).

4. Kobayashi, T. & Nishizawa, N. K. Iron Uptake, Translocation, and Regulation in Higher Plants. Annu. Rev. Plant Biol. 63, 131–152 (2012).

5. Nozoye, T. et al. Phytosiderophore Efflux Transporters Are Crucial for Iron Acquisition in Graminaceous Plants. J. Biol. Chem. 286, 5446–5454 (2011).

6. Curie, C., Panaviene, Z., Loulergue, C. & Dellaporta, S. L. Maize yellow stripe1 encodes a membrane protein directly involved in Fe (III) uptake. Nature (2001).

7. Murata, Y. et al. A specific transporter for iron(III)-phytosiderophore in barley roots. The Plant Journal 46, 563–572 (2006).

8. Takagi, S.-I. Naturally occurring iron-chelating compounds in oat- and rice-root washings. Soil Science and Plant Nutrition 22, 423–433 (1976).

9. Santi, S. & Schmidt, W. Dissecting iron deficiency-induced proton extrusion in Arabidopsis roots. New Phytologist 183, 1072–1084 (2009).

10. Robinson, N. J., Procter, C. M., Connolly, E. L. & Guerinot, M. L. A ferric-chelate reductase for iron uptake from soils. Nature (1999).

11. Eide, D., Broderius, M., Fett, J. & Guerinot, M. L. A novel iron-regulated metal transporter from plants identified by functional expression in yeast. Proceedings of the National Academy of Sciences 93, 5624–5628 (1996).

12. Connolly, E. L. Expression of the IRT1 Metal Transporter Is Controlled by Metals at the Levels of Transcript and Protein Accumulation. THE PLANT CELL ONLINE 14, 1347–1357 (2002).

13. Romheld, V. & Marschner, H. Evidence for a Specific Uptake System for Iron Phytosiderophores in Roots of Grasses. PLANT PHYSIOLOGY 80, 175–180 (1986).

14. Lan, P. et al. iTRAQ Protein Profile Analysis of Arabidopsis Roots Reveals New Aspects Critical for Iron Homeostasis. PLANT PHYSIOLOGY 155, 821–834 (2011).

15. Rodriguez-Celma, J. et al. Mutually Exclusive Alterations in Secondary Metabolism Are Critical for the Uptake of Insoluble Iron Compounds by Arabidopsis and Medicago truncatula. PLANT PHYSIOLOGY 162, 1473–1485 (2013).

16. Jin, C. W. et al. Iron Deficiency-Induced Secretion of Phenolics Facilitates the Reutilization of Root Apoplastic Iron in Red Clover. PLANT PHYSIOLOGY 144, 278–285 (2007).

17. Sisó-Terraza, P., Rios, J. J., Abadía, J., Abadía, A. & Álvarez-Fernández, A. Flavins secreted by roots of iron-deficient Beta vulgarisenable mining of ferric oxide via reductive mechanisms. New Phytol 209, 733–745 (2015).

18. Fourcroy, P. et al. Involvement of the ABCG37 transporter in secretion of scopoletin and derivatives by Arabidopsisroots in response to iron deficiency. New Phytol 201, 155–167 (2013).

19. Schmid, N. B. et al. Feruloyl-CoA 6’-Hydroxylase1-Dependent Coumarins Mediate Iron Acquisition from Alkaline Substrates in Arabidopsis. PLANT PHYSIOLOGY 164, 160–172 (2014).

20. Schmidt, H. et al. Metabolome Analysis of Arabidopsis thaliana Roots Identifies a Key Metabolic Pathway for Iron Acquisition. PLoS ONE 9, e102444 (2014).

21. Kai, K. et al. Scopoletin is biosynthesized via ortho-hydroxylation of feruloyl CoA by a 2-oxoglutarate-dependent dioxygenase in Arabidopsis thaliana. The Plant Journal 55, 989–999 (2008).

22. Mladěnka, P. et al. In vitro interactions of coumarins with iron. Biochimie 92, 1108–1114 (2010).

23. Murgia, I., Tarantino, D., Soave, C. & Morandini, P. Arabidopsis CYP82C4 expression is dependent on Fe availability and circadian rhythm, and correlates with genes involved in the early Fe deficiency response. Journal of Plant Physiology 168, 894–902 (2011).

24. Kruse, T. et al. In Planta Biocatalysis Screen of P450s Identifies 8-Methoxypsoralen as a Substrate for the CYP82C Subfamily, Yielding Original Chemical Structures. Chemistry & Biology 15, 149–156 (2008).

25. Ray, A. B. et al. Structures of cleomiscosins, coumarinolignoids of cleome viscosa seeds. Tetrahedron 41, 209–214 (1985).

26. Yuan, C. et al. Metal-free oxidation of aromatic carbon–hydrogen bonds through a reverse-rebound mechanism. Nature 499, 192–196 (2013).

27. Fujita, P. A. et al. The UCSC Genome Browser database: update 2011. Nucleic Acids Research 39, D876–D882 (2010).

28. Kanehisa, M. KEGG: Kyoto Encyclopedia of Genes and Genomes. Nucleic Acids Research 28, 27–30 (2000).

29. Amtmann, A. Abiotic Stress and Plant Genome Evolution. Search for New Models. PLANT PHYSIOLOGY 138, 127–130 (2005).

30. Smith, S. A., Beaulieu, J. M. & Donoghue, M. J. An uncorrelated relaxed-clock analysis suggests an earlier origin for flowering plants. Proceedings of the National Academy of Sciences 107, 5897–5902 (2010).

31. Susín, S. et al. Flavin excretion from roots of iron-deficient sugar beet (Beta vulgaris L.). Planta 193, 514–519 (1994).

32. Peters, N. K. Current ReviewPhenolic Compounds as Regulators of Gene Expression in Plant-Microbe Interactions. MPMI 3, 4 (1990).

33. Brown, J. C., Chaney, R. L. & Ambler, J. E. A New Tomato Mutant Inefficient in the Transport of Iron. Physiol Plant 25, 48–53 (1971).

34. Brown, J. C. & Ambler, J. E. ‘Reductants’ Released by Roots of Fe-Deficient Soybeans1. Agronomy Journal 65, 311 (1973).

35. Sisó-Terraza, P. et al. Accumulation and Secretion of Coumarinolignans and other Coumarins in Arabidopsis thaliana Roots in Response to Iron Deficiency at High pH. Front. Plant Sci. 7, 327 (2016).

36. Brutinel, E. D. & Gralnick, J. A. in Microbial Metal Respiration 83–105 (Springer Berlin Heidelberg, 2012). doi:10.1007/978-3-642-32867-1_4

37. Price-Whelan, A., Dietrich, L. E. P. & Newman, D. K. Rethinking ‘secondary’ metabolism: physiological roles for phenazine antibiotics. Nat Chem Biol 2, 71–78 (2006).

